# Missense and nonsense mutations of the zebrafish *hcfc1a* gene result in contrasting mTor and radial glial phenotypes

**DOI:** 10.1101/2022.10.21.513292

**Authors:** Victoria L. Castro, David Paz, Valeria Virrueta, Igor L. Estevao, Brian I. Grajeda, Cameron C. Ellis, Anita M. Quintana

## Abstract

Mutations in the HCFC1 transcriptional co-factor protein are the cause of *cblX* syndrome and X-linked intellectual disability (XLID). *cblX* is the more severe disorder associated with intractable epilepsy, abnormal cobalamin metabolism, facial dysmorphia, cortical gyral malformations, and intellectual disability. *In vitro*, *Hcfc1* regulates neural precursor (NPCs) proliferation and number, which has been validated in zebrafish. However, conditional deletion of *Hcfc1* in Nkx2.1+ NPCs increased cell death, reduced *Gfap* expression, and reduced numbers of GABAergic neurons. Thus, the role of HCFC1 in brain development is not completely understood. Recently, knock-in of both a *cblX* (*HCFC1*) and *cblX*-like (*THAP11*) allele were created in mice. Knock-in of the *cblX*-like allele was associated with increased expression of proteins required for ribosome biogenesis. However, the brain phenotypes were not comprehensively studied due to sub-viability and therefore, a mechanism underlying increased ribosome biogenesis was not described. We used a missense, a nonsense, and two conditional zebrafish alleles to further elucidate this mechanism during brain development. We observed contrasting phenotypes at the level of Akt/mTor activation, the number of radial glial cells, and the expression of two downstream target genes of HCFC1, *asxl1* and *ywhab*. Despite these divergent phenotypes, each allele studied demonstrates with a high degree of face validity when compared to the phenotypes reported in the literature. Collectively, these data suggest that individual mutations in the HCFC1 protein result in differential mTor activity which is associated with contrasting cellular phenotypes.

## Introduction

Mutations in the HCFC1 transcriptional co-factor protein cause *cblX* syndrome (MIM 309541) (Gérard et al., 2015, 1; Yu et al., 2013) and X-linked intellectual disability (XLID) (Huang et al., 2012; Jolly et al., 2015; Koufaris et al., 2016a, 1; Piton et al., 2011; Piton et al., 2013). *cblX* is a multiple congenital anomaly syndrome characterized by abnormal cobalamin metabolism, neurodevelopmental defects, cortical gyral malformations, intractable epilepsy, craniofacial dysmorphic features, and movement disorders. However, XLID is milder and associated only with intellectual disability. The clinical phenotypes of the patients are heterogeneous and varied in severity. Multiple systems and approaches have been undertaken to understand the function of HCFC1 during development. *In vitro* over expression assays first revealed decreased neural precursor cell (NPCs) proliferation with abnormal differentiation and a bias towards the astrocyte lineage in mouse neurospheres (Huang et al., 2012). Later we demonstrated that mutation or knockdown of the zebrafish paralogs of *HCFC1* caused increased proliferation of NPCs, but these changes were associated with increased expression of radial glial (RGC) and neuronal markers (Castro et al., 2020; Quintana et al., 2017). Global deletion of *Hcfc1* in mice was embryonic lethal (Minocha et al., 2016a), but tissue specific deletion in Nkx2.1+ progenitors caused increased NPC death and a decrease in GABAergic interneurons and glia (Minocha and Herr, 2019). Thus, while model system specific, previous studies collectively demonstrate a function for HCFC1 in brain development and neural precursors.

Recently, a patient derived *cblX* mutation (p.A115V) was knocked into the mouse *Hcfc1* loci to produce the first *in vivo cblX* model, but the brain phenotypes were not comprehensively studied. However, these studies suggested a mechanism by which mutations in HCFC1 cause abnormal development. Chern and colleagues uncovered increased expression of proteins essential for ribosome biogenesis in a *cblX*-like disorder (Chern et al., 2022) caused by mutations in the THAP11 (p.F80L) protein, which interacts with HCFC1 (Mazars et al., 2010). These data are supported by previous studies which have shown that HCFC1 regulates genes essential for metabolism and stem cell maintenance (Dejosez et al., 2008; Dejosez et al., 2010). What remains uncharacterized is the mechanism by which mutations in the HCFC1 protein promote increased ribosome biogenesis and the effects of this biogenesis on brain development.

The mechanistic target of rapamycin (mTor) pathway is a well-known regulator of protein translation and cell growth, particularly as it relates to stem cell regulation (Gabut et al., 2020). Interestingly, the mTor pathway is a common pathway dysregulated in neurodevelopmental disorders (Parenti et al., 2020). mTor is regulated at many levels, but one example of a positive regulator of mTor is the PI3K/Akt pathway. Activation of PI3K leads to Akt activation, which in turn drives activation of mTor signaling and promotes ribosomal biogenesis and protein synthesis. We have previously demonstrated that inhibition of PI3K can restore the NPC deficits present in a zebrafish with a nonsense mutation in *hcfc1a* (co60 allele) (Castro et al., 2020). These data, combined with recent literature, indirectly link regulation of PI3K/Akt signaling with phenotypes that occur in *cblX* syndrome. In addition, we identified a driver of Akt signaling, the Asxl1 polycomb group protein, as up-regulated in the co60 allele implicating this HCFC1 downstream target gene as one putative mediator of Akt/mTor in *cblX*. Therefore, we hypothesized that mutations in HCFC1 drive ribosomal biogenesis and disrupt brain development through regulation of Akt/mTor.

Here we identified divergent activity of the Akt/mTor signaling pathway in two independent zebrafish *hcfc1a* germline mutant alleles. Contrasting Akt/mTor activity was correlated with divergent RGC phenotypes. In addition, we identified disparate expression of two downstream effectors of HCFC1 in each allele: Asxl1 and 14-3-3βα, both of which are known to regulate Akt activity and brain development (An et al., 2019, 1; Cornell and Toyo-oka, 2017; Gómez-Suárez et al., 2016a; Youn et al., 2017). Further, restoration experiments revealed that *gfap* expression is mTor dependent. Thus, our analysis of distinct *hcfc1a* mutant alleles suggests that unique variants can have different effects on HCFC1 protein dosage/function and can lead to individual cellular and molecular phenotypes.

## Results

### Akt/mTOR signaling is increased in the co60 allele

We have previously established that nonsense mutation of *hcfc1a* (co60 allele) causes an increase in *asxl1* expression, which is associated with increased proliferation of NPCs (Castro et al., 2020). Asxl1 promotes Akt phosphorylation (Youn et al., 2017) and we established that inhibition of PI3K, an upstream regulator of Akt, is sufficient to restore NPC numbers to normal levels in the co60 allele. Thus, we hypothesized that nonsense mutation of *hcfc1a* leads to increased activation of Akt kinase. We performed western blot analysis using anti-phosho-Akt threonine 308 (pAkt308) and total Akt antibodies. We detected increased expression of total Akt, which was associated with hyperphosphorylation of pAkt308 (Fig. 1A). Since total Akt was routinely higher in the co60 allele, we used β-actin as an additional loading control to validate increased expression of total Akt protein (Fig 1A). We observed equal β-actin expression, which correlated with Ponceau S staining of the membrane. We did not analyze phosphorylation of Serine 473, as we could not identify an antibody that would cross react with zebrafish Akt at this location.

**Figure 1:**
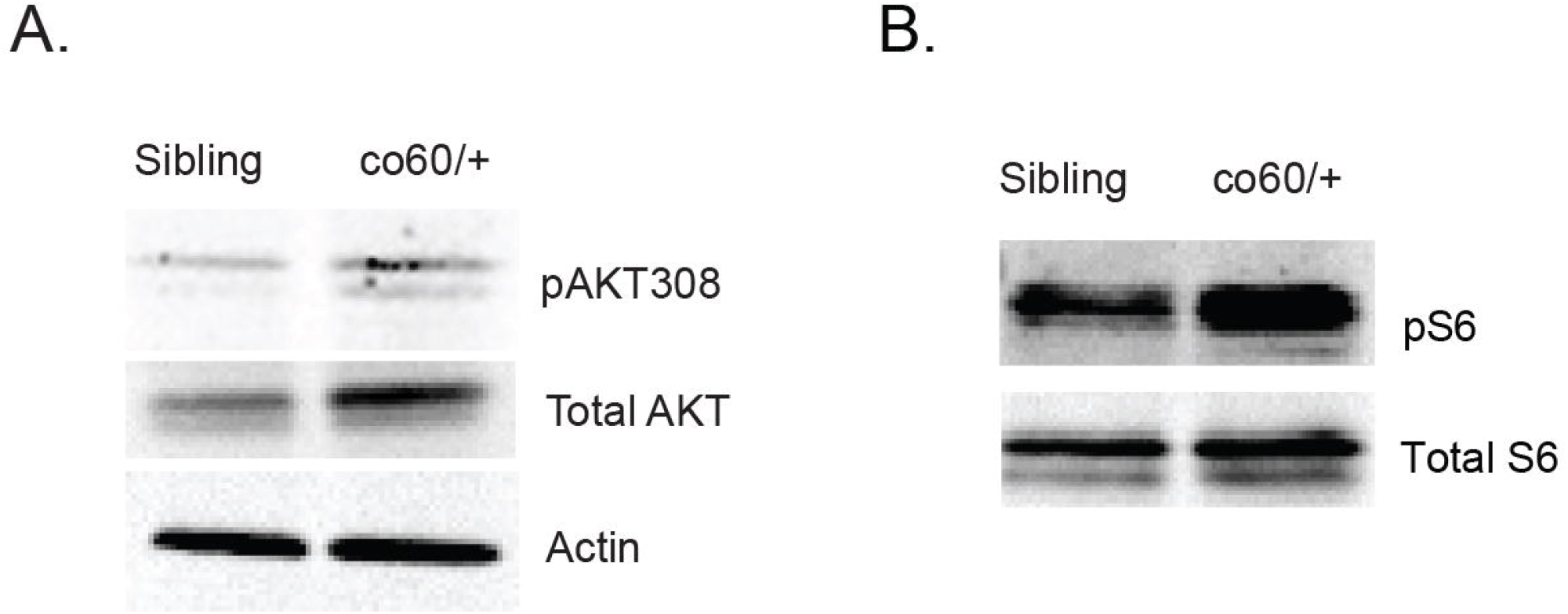
Akt/mTor signaling is increased in the *hcfc1a^co60/+^* allele. A-B. Western blot analysis was performed with anti-phospho Akt (thr308) antibodies (pAKT308), total Akt, anti-phospho p70S6kinase (pS6),total S6, or β-actin using brain homogenates from sibling wildtype or carriers of the *hcfc1a^co60/+^* (co60/+) allele. N=4/group/biological replicate and performed in 2 independent biological replicates.

Akt kinase is an upstream regulator of the mTor pathway, which subsequently promotes growth, proliferation, translation, and ribosomal protein synthesis. Chern and colleagues published that *cblX* is associated with increased expression of proteins required for ribosome biogenesis (Chern et al., 2022). Therefore, we performed western blot analysis to detect phosphorylation of S6 kinase (pS6), which is phosphorylated by active mTor and represents a marker of mTor activity. We observed increased pS6 kinase in carriers of the co60 allele with equivalent levels of total S6 protein (Fig 1B). We did not use β-actin to validate protein loading as total S6 has been used as a proper loading control in similar assay (Ventura et al., 2010). Ponceau S was used to confirm the total S6. In addition, we did not detect differences in total S6 in any biological replicate. These data demonstrate that nonsense mutation of *hcfc1a* is associated with increased Akt/mTor signaling.

### Missense mutation of the N-terminal kelch domain does not change expression of *hcfc1a*

Nonsense mutation of *hcfc1a* results in increased Akt/mTor (Fig 1). However, nonsense mutation of *hcfc1a* (co60 allele) is haploinsufficient in zebrafish and reduces *hcfc1a* expression by 50%, which does not accurately reflect the types of mutations present in *cblX* syndrome. *cblX* syndrome is caused by missense mutations that do not reduce overall protein expression (Yu et al., 2013). We generated an additional *hcfc1a* allele (*hcfc1a^co64^* or co64) which results in an insertion deletion that causes an in-frame deletion of arginine at amino acid position 75 with two missense mutations p.G76M and p.D77N within the kelch protein interaction domain (Fig 2A). The co64 allele is homozygous viable into adulthood (the only model to date) and homozygous offspring obey Mendelian inheritance patterns. As shown in Fig 2B, we did not detect any significant difference in the expression of *hcfc1a* or *hcfc1b*, a second zebrafish ortholog of *HCFC1* in the co64 allele.

**Figure 2:**
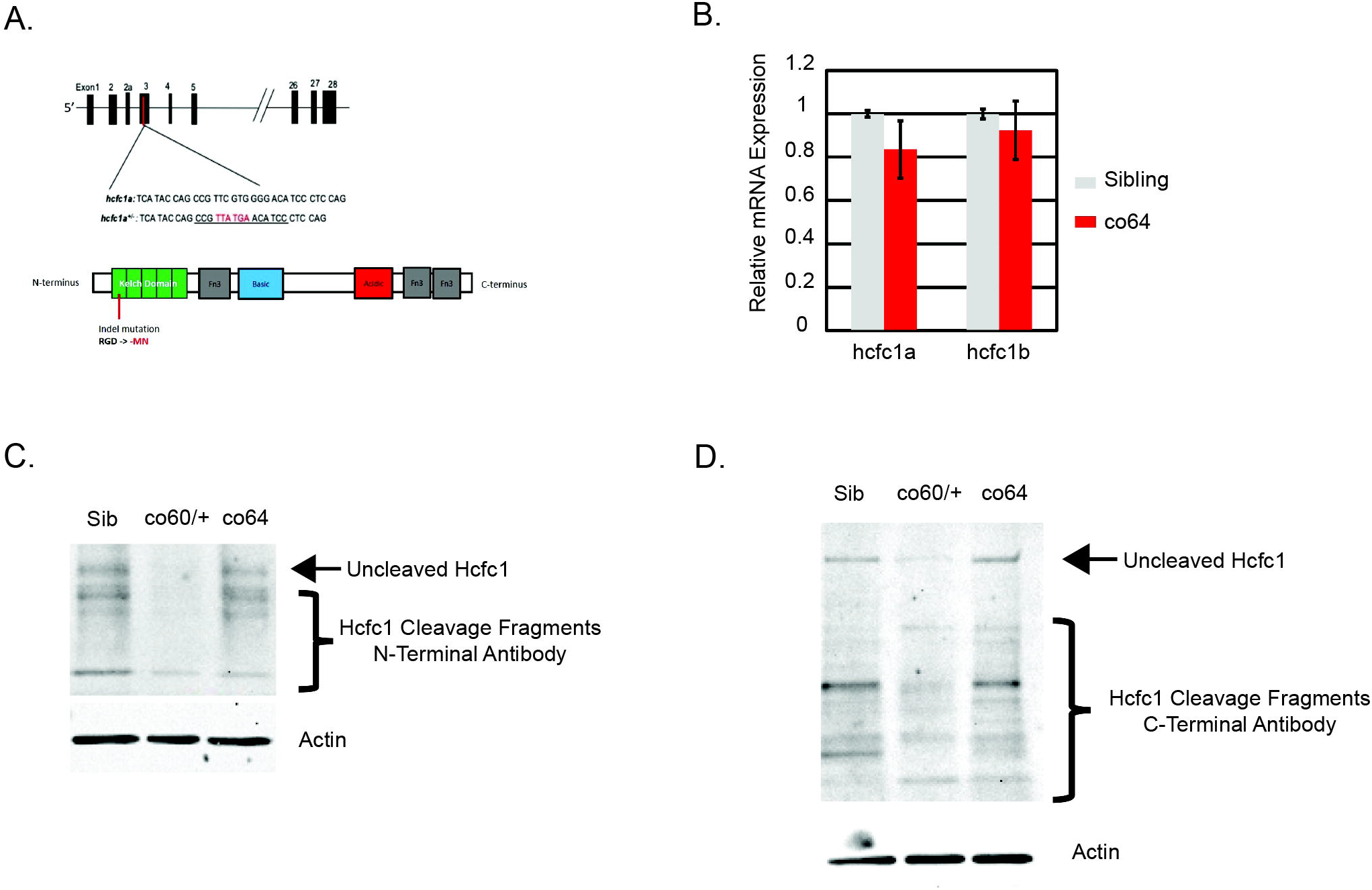
Missense and nonsense alleles of the *hcfc1a* gene differentially affect Hcfc1 expression. A. Schematic diagram for the location of the *hcfc1a^co64^* homozygous viable allele. CRISPR/Cas9 created and insertion/deletion in the genomic DNA that causes an in-frame deletion of arginine (R) and a 2-amino acid base pair change (GD>MN) in the kelch domain. B. Quantitative PCR (qPCR) was performed with total RNA isolated from whole brain homogenates of wildtype siblings or homozygous carriers of the *hcfc1a^co64^* allele (co64). The expression of *hcfc1a* and the *hcfc1b* paralog were analyzed. No significant difference in expression was observed, consistent with the effects of missense mutations in *cblX* syndrome. A total of N=23 wildtype sibling and N=25 homozygous brains were used to measure *hcfc1a* expression. Expression of *hcfc1b* was analyzed with a total number of N=18 wildtype siblings and N=20 homozygous carriers. Numbers were obtained over the course of a minimum of 3 biological replicates. C-D. Western blot analysis was performed with N-terminal (C) or C-terminal (D) specific antibodies to the HCFC1 protein. Total uncleaved and cleaved protein was detected in brain homogenates from wildtype siblings, heterozygous carriers of the *hcfc1a^co60/+^* (co60/+), and homozygous carriers of the *hcfc1a^co64^* (co64) allele. β-actin was used as a loading control. N=4/group/biological replicate and performed in 2 independent biological replicates were utilized for western blot analysis.

### The co60 and co64 alleles have unique effects on Hcfc1a protein expression

HCFC1 is synthesized as a large preprotein that is cleaved into N and C-terminal fragments (Kapuria et al., 2016). Consequently, nonsense mutations that cause premature stop codons early in the protein sequence are likely to have different effects on the expression of all cleavage fragments produced whereas missense mutations are hypothesized to affect only a single domain. We compared the expression of both N and C-terminal fragments in the co60 and co64 alleles by western blot using antibodies that independently detect the N or C-terminal proteolytical cleavage fragments. The co60 allele (nonsense) decreased expression of N- and C-terminal fragments (Fig 2C&D). In contrast, the co64 allele (missense) did not markedly reduce overall protein expression of either fragment, which indicated that the N-terminal missense fragment was produced at close to the same level relative to sibling wildtype (Fig 2C&D). These data also provide evidence that the co64 allele does not reduce overall protein expression, but the co60 allele affects the expression level of Hcfc1a.

### The co64 allele is associated with decreased Akt/mTor signaling

We observed increased Akt/mTor in the co60 allele, but since the co64 and co60 alleles differentially affect protein expression, we hypothesized that dosage changes in HCFC1 expression (co60) would affect brain development and Akt/mTor signaling differently than missense mutation in the kelch domain. We used western blot analysis to determine the level of Akt/mTor signaling in the co64 allele. We observed hypophosphorylation of Akt at threonine 308 with equivalent levels of total Akt (Fig 3A). We used total Akt as a loading control as we did not detect changes in the expression of total Akt in any of the biological replicates performed. This contrasts with the co60 allele, which demonstrated consistently with increased total Akt. Decreased Akt phosphorylation was associated with decreased pS6 (Fig 3B) and total S6 protein was unchanged across multiple biological replicates. These data reveal differential regulation of Akt/mTor in the co60 and co64 alleles, which we confirmed have unique effects on Hcfc1a protein expression (Fig. 2).

**Figure 3:**
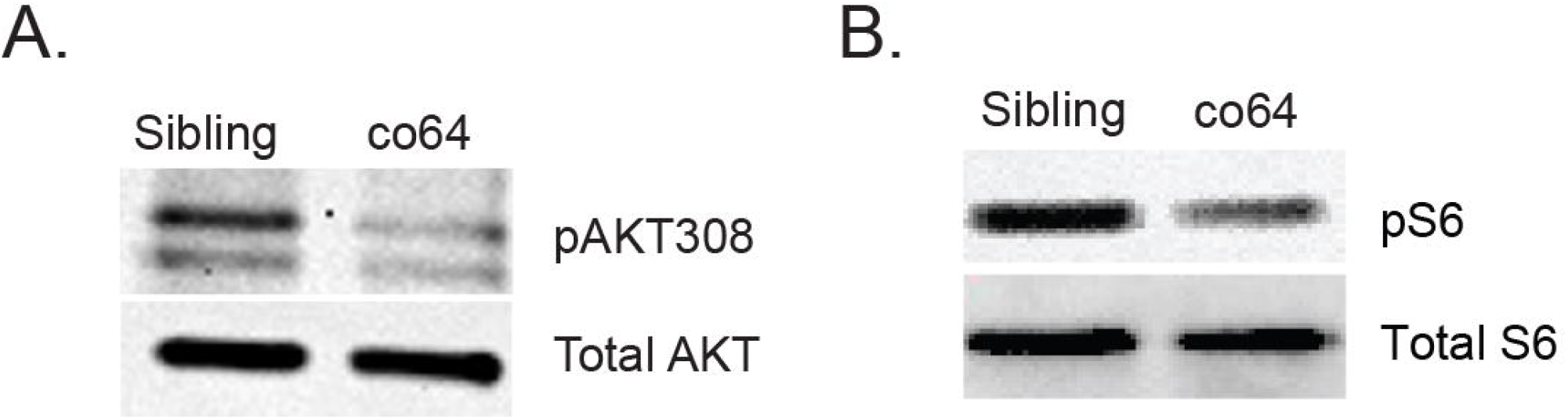
Differential regulation of Akt/mTor in the *hcfc1a^co64^* allele. A-B. Western blot analysis was performed with anti-phospho Akt (thr308) (pAKT308), total Akt, anti-phospho-p70S6kinase (pS6),or total S6 antibodies using brain homogenates from sibling wildtype or carriers of the *hcfc1a^co64^*(co64) allele. N=4/group/biological replicate and performed in 2 independent biological replicates were utilized for western blot analysis.

### NPC number is increased in *cblX*-like syndromes

We have previously demonstrated that the co60 allele causes increased numbers of Sox2+ NPCs (Castro et al., 2020). We sought to determine the number of NPCs in the co64 allele given the differential regulation of Akt/mTor. To quantify the number of NPCs, we first crossed the co64 allele into the *Tg*(*sox2:2A:EGFP*) reporter. We then produced a single cell homogenate and preformed flow cytometry to detect EGFP positive cells. We detected an increase in the number of GFP+ cells by flow cytometry at 6 days post fertilization (DPF) (Fig 4A&A’) and validated this increase using immunohistochemistry at 2 DPF (Figure 4B&B’, arrows indicate regions of increased cell number). We performed flow cytometry on two independent occasions and found a statistically significant (p<0.05) increase in NPCs in heterozygous and homozygous carriers of the co64 allele (Fig 4C). These data suggest that both the co60 and co64 allele have increased NPCs. The presence of overlapping phenotypes provides face validity for the effectiveness of each allele as a putative model system to inform as to the function of HCFC1 in NPC development.

**Figure 4:**
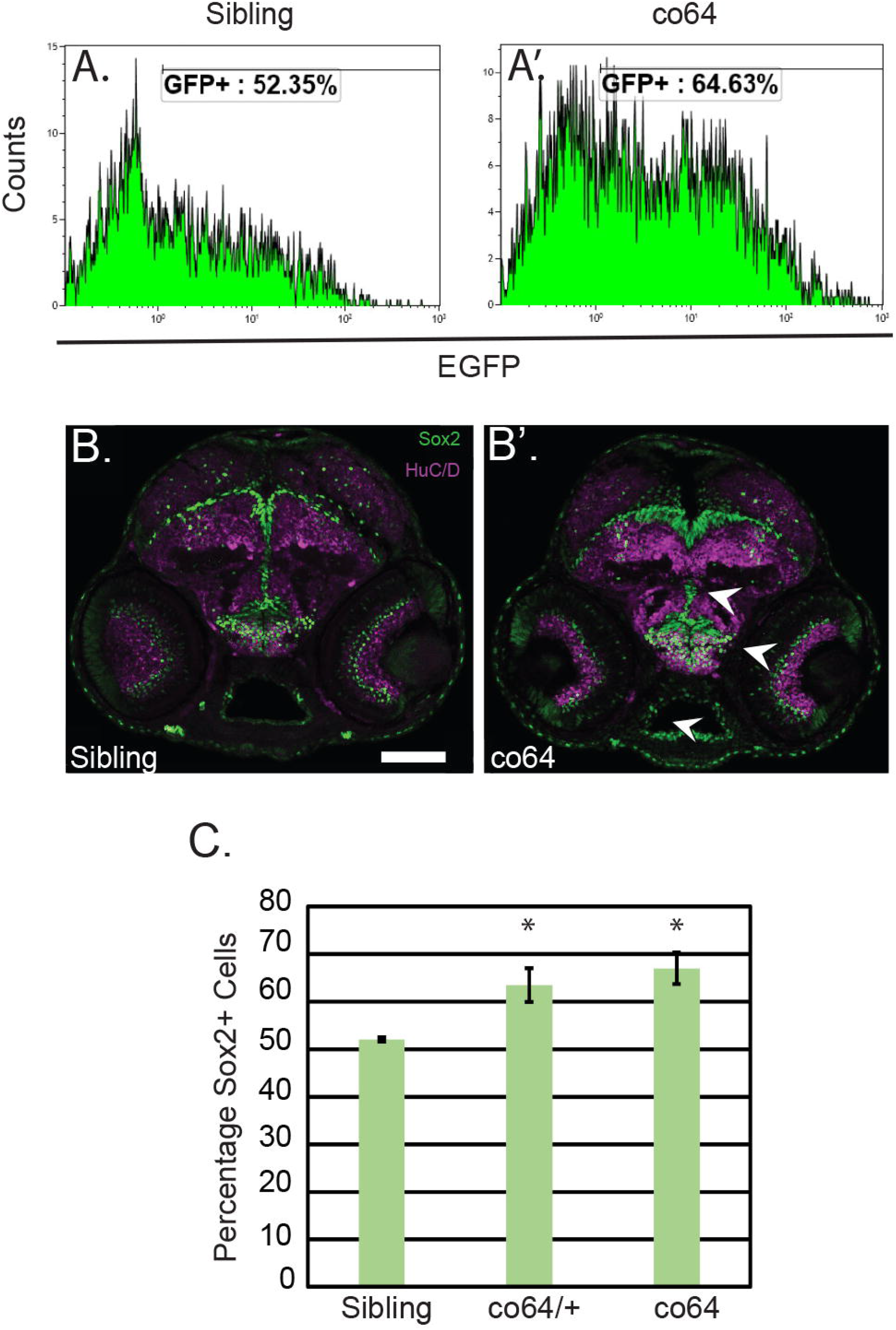
Neural precursors (NPCs) are increased in the co64 allele. A&A’. Flow cytometry analysis was used to detect the number of Sox2+ cells in the *hcfc1a^co64^* allele (co64) and wildtype siblings at 6 days post fertilization (DPF). Histograms represent a single biological replicate with N=5/group. B&B’. Immunohistochemistry at 2 days post fertilization (DPF) was performed to detect Sox2+ (green) and Huc/D+ cells (magenta) in sibling wildtype and the co64 allele. N=6/group obtained from 2 biological replicates. C. Quantification of percentages obtained by flow cytometry from 3 biological replicates (N=5/replicate and performed in 3 biological replicates, one is shown in A&A’). The percentage of Sox2+ cells was quantified by flow cytometry and the average of 3 replicates is plotted. *p<0.05.

### The co60 and co64 alleles differentially disrupt RGC number

We have previously shown that the co60 allele increases the expression of *gfap* and *elavl3*, markers of RGCs and neurons, respectively (Castro et al., 2020). Therefore, we measured the expression of both genes in the co64 allele. We observed a statistically significant decrease in *gfap* expression at 2 DPF (Fig 5C), which we validated by immunohistochemistry at 6 DPF (Fig 5A&B). We next crossed the co64 allele into the *Tg*(*gfap:EGFP*) reporter and performed flow cytometry to quantify the total number of RGCs. We observed a statistically significant decrease in the total number of RGCs (Fig 5D). Decreased RGC number in the co64 allele contrasted with the number of RGCs in the co60 allele, as we detected equivalent numbers of RGCs using flow cytometry in the co60 allele (Fig 5E). Interestingly, we validated an increase in the expression of *gfap* in the co60 allele (Fig 6E, yellow bars, p=0.06). These data demonstrate that 1) the co64 allele reduces the number of RGCs, 2) the co60 allele increases *gfap* expression, but has no effect on total RGC number, and 3) the different *hcfc1a* alleles studied have unique effects on RGC development.

**Figure 5:**
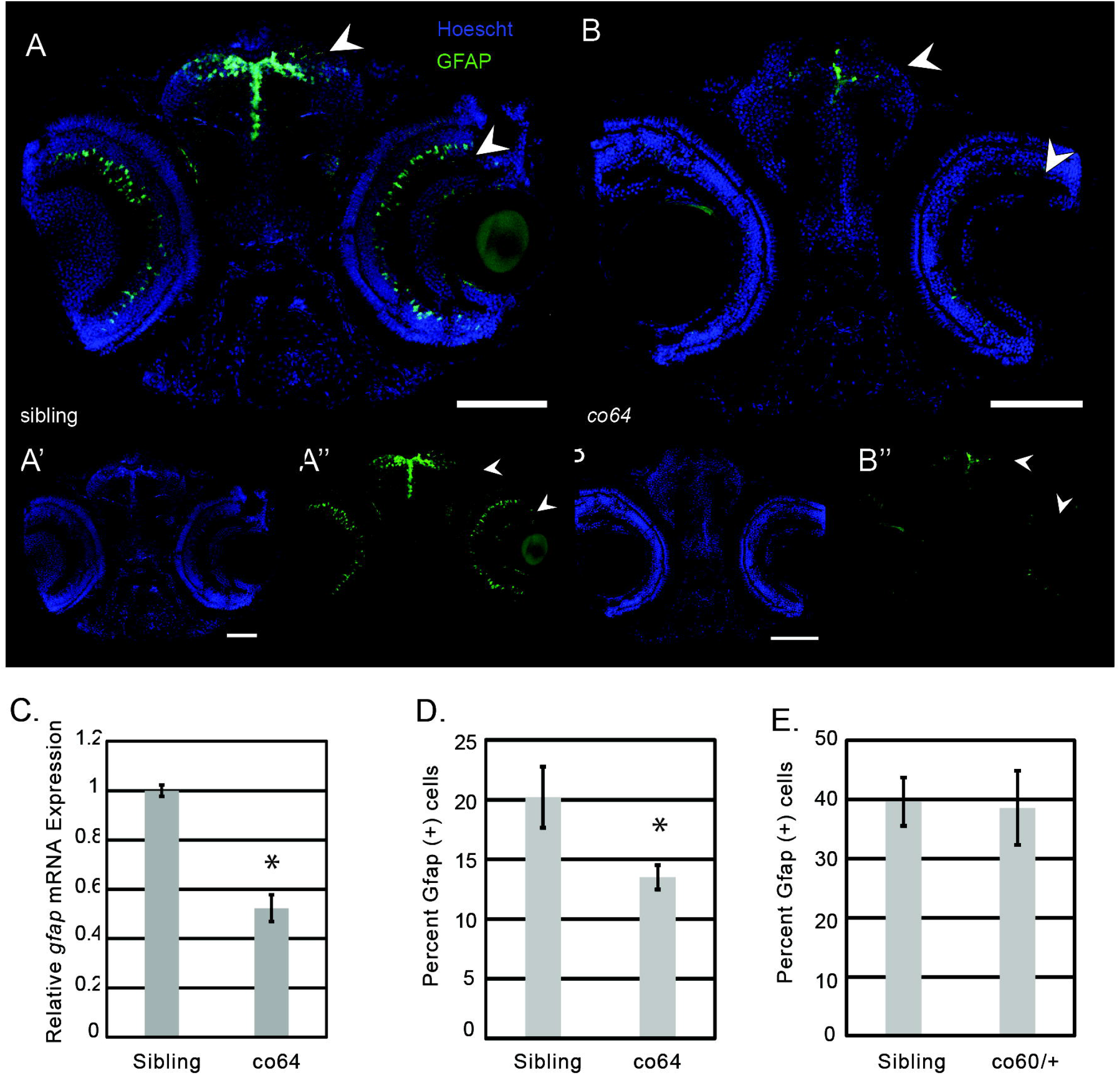
Radial glial cells (RGCs) are reduced in number and expression in the co64 allele. A-B. A’-B’, & A”-B”. Immunohistochemistry was used to detect the Gfap+ cells in wildtype sibling and homozygous carriers of the *hcfc1a^co64^* allele (co64) at 6 days post fertilization (DPF). The co64 allele was crossed with the *Tg*(*gfap:EGFP*) reporter. Hoescht DNA content stain (A’-B’) was used as a control. N=4 sibling and N=7 co64. Arrows indicate areas of decreased expression. C. Quantification of *gfap* expression in sibling wildtype and homozygous carriers of the co64 allele. Total RNA was isolated from whole brain homogenates and *gfap* expression was quantified by qPCR from 3 biological replicates (N=3/replicate) (*p<0.05). D. The number of Gfap+ cells was quantified at 5 DPF using flow cytometry in the co64 allele and wildtype sibling. Analysis was performed in biological triplicate with a total N=14 and the average percentage of positive cells was plotted (p<0.05). E. The number of Gfap+ cells was quantified by flow cytometry in sibling wildtype and the *hcfc1a^co60^* allele. Two biological replicates were performed with total N=6. The average percentage of cells in each biological replicate is plotted with error bars representing standard error of the mean between biological replicates. No significant difference was found.

**Figure 6:**
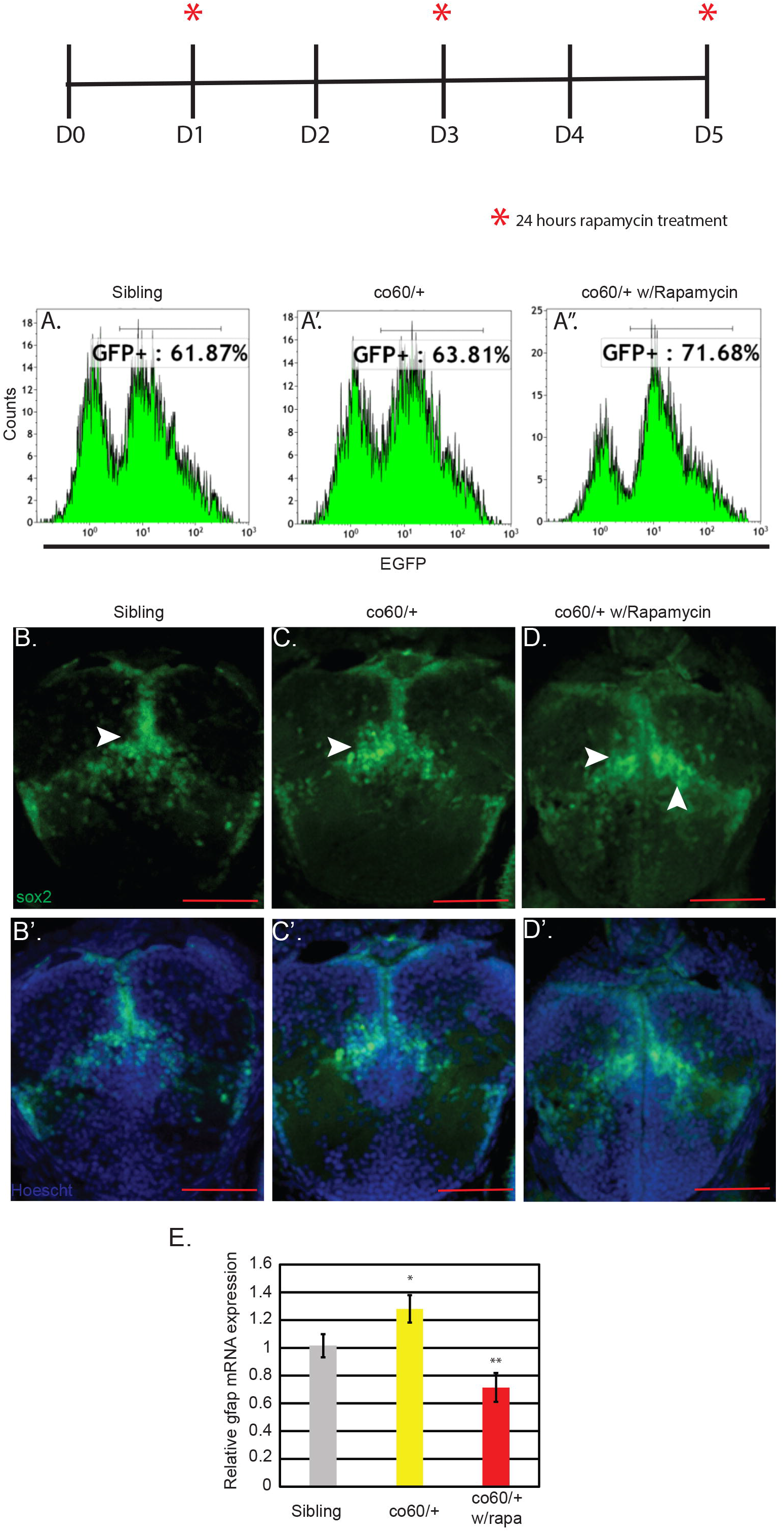
*gfap* expression but not Sox2 cell number is mTor dependent. Top: Schematic representing the onset of rapamycin treatment. Each asterisk represents the onset of a 24- hour period of treatment. Days (D) without an asterisk are those where media did not contain rapamycin.A-A”. Flow cytometry of the *hcfc1a^co60/+^(*co60/+) and wildtype sibling was performed at 5 days post fertilization (DPF). The *hcfc1a^co60/+^* allele was treated with rapamycin in an independent sample to inhibit mTor signaling (w/rapamycin). The number of Sox2+ cells were quantified with the *Tg*(*sox2:2A:EGFP*) transgene. N=2 heads per group (WT, Co60, Co60 w/rapamycin). The number of Sox2+ cells was not restored by treatment with rapamycin. B-D. Immunohistochemistry validation of A-A” analyzing Sox2+ using the *Tg*(*sox2:2A:EGFP*). Hoescht stain was performed as a control in B’-D’. E. Quantitative PCR (qPCR) was used to determine the expression of *gfap* in sibling wildtype, the co60/+ allele, or the co60/+ treated with rapamycin (w/rapa). The level of *gfap* expression is increased in the co60/+ allele (*p<0.06) and restored by treatment with rapamycin (**p<0.05).

### NPC number is mTor independent

Both the co60 and co64 alleles cause an increase in the number of NPCs but demonstrate divergent regulation of Akt/mTor. Based on these data, we surmised that NPC number was Akt/mTor independent. To test this, we treated co60 embryos/larvae with rapamycin (Fig 6 Schematic) and measured the number of Sox2+ NPCs by flow cytometry. We consistently observed a 3-5% overall increase in the number of NPCs by flow cytometry in the co60 allele (Fig 6A&A’). These data are consistent with previous studies (Castro et al., 2020). Treatment of rapamycin did not reduce the number of NPCs at 5 DPF (Fig. 6A-A”) but instead exacerbated the phenotype leading to an even higher number of total EGFP Sox2+ cells. We validated these results with immunohistochemistry (Fig 6B-D).

### *gfap* expression is partially regulated by mTor in the co60 allele

The co60 and co64 alleles demonstrated with differential expression of *gfap* as documented in Fig 5C and (Castro et al., 2020). In the co60 allele, increased *gfap* expression is not due to increased numbers of cells (Fig 5E). We validated the increased expression of *gfap* expression in the co60 allele in Fig 6E (yellow bars, p=0.06). Since *gfap* and mTor activity were differentially regulated in the co60 and co64, we sought to determine if *gfap* expression was mTor dependent. Wildtype and co60 carriers were treated with rapamycin as shown in Fig 6 schematic and the expression of *gfap* was analyzed by qPCR analysis. Treatment with rapamycin caused a statistically significant decrease in the expression of *gfap* in mutant animals (co60) (Fig 6E), red bars). These data suggest that *gfap* expression is partially regulated by mTor signaling in the co60 allele.

### *Asxl1* expression correlates with Akt/mTor activity in the co60 and co64 alleles

We have previously established that the co60 allele caused increased expression of *asxl1*, a mediator of Akt signaling (Castro et al., 2020; Youn et al., 2017). Thus, we hypothesized that differential expression of *asxl1* was mediating contrasting Akt/mTor signature in each allele. We measured the expression of *asxl1* in the co64 and observed it to be significantly decreased relative to sibling wildtype (Fig 7A). Human HCFC1 protein has been shown to bind to the *ASXL1* promoter (Dehaene et al., 2020) and therefore, we hypothesized that the expression of *asxl1* was reduced in the co64 allele because missense mutations disrupt binding to the *ASXL1* promoter. To test these this hypothesis, we utilized the *Tg*(*hsp701:HCFC1*) transgene (Castro et al., 2020). After performing a heat shock protocol (Castro et al., 2020; Hudish et al., 2013), we performed chromatin immunoprecipitation (ChIP) and used PCR to detect binding of wildtype human HCFC1 to the zebrafish *mmachc* and *asxl1* promoters. *Mmachc* is a known downstream target gene of HCFC1 and used here as a positive control (Dejosez et al., 2010). Semi-quantitative PCR was performed in technical triplicate and demonstrated amplification of *mmachc* and *asxl1* promoter regions in ChIP assays performed with HCFC1 specific antibodies (Fig 7B). We did not detect amplification after 40 cycles in the IgG antibody control lanes, except in one sample designed to amplify the *asxl1* promoter region. However, this amplification was far less than the enrichment detected using HCFC1 specific antibodies. These results indicate putative binding to the *mmachc* and *asxl1* promoters by human HCFC1. Positive binding was associated with increased mRNA expression of *mmachc* and *asxl1* after heat shock protocol (Fig 7C).

**Figure 7:**
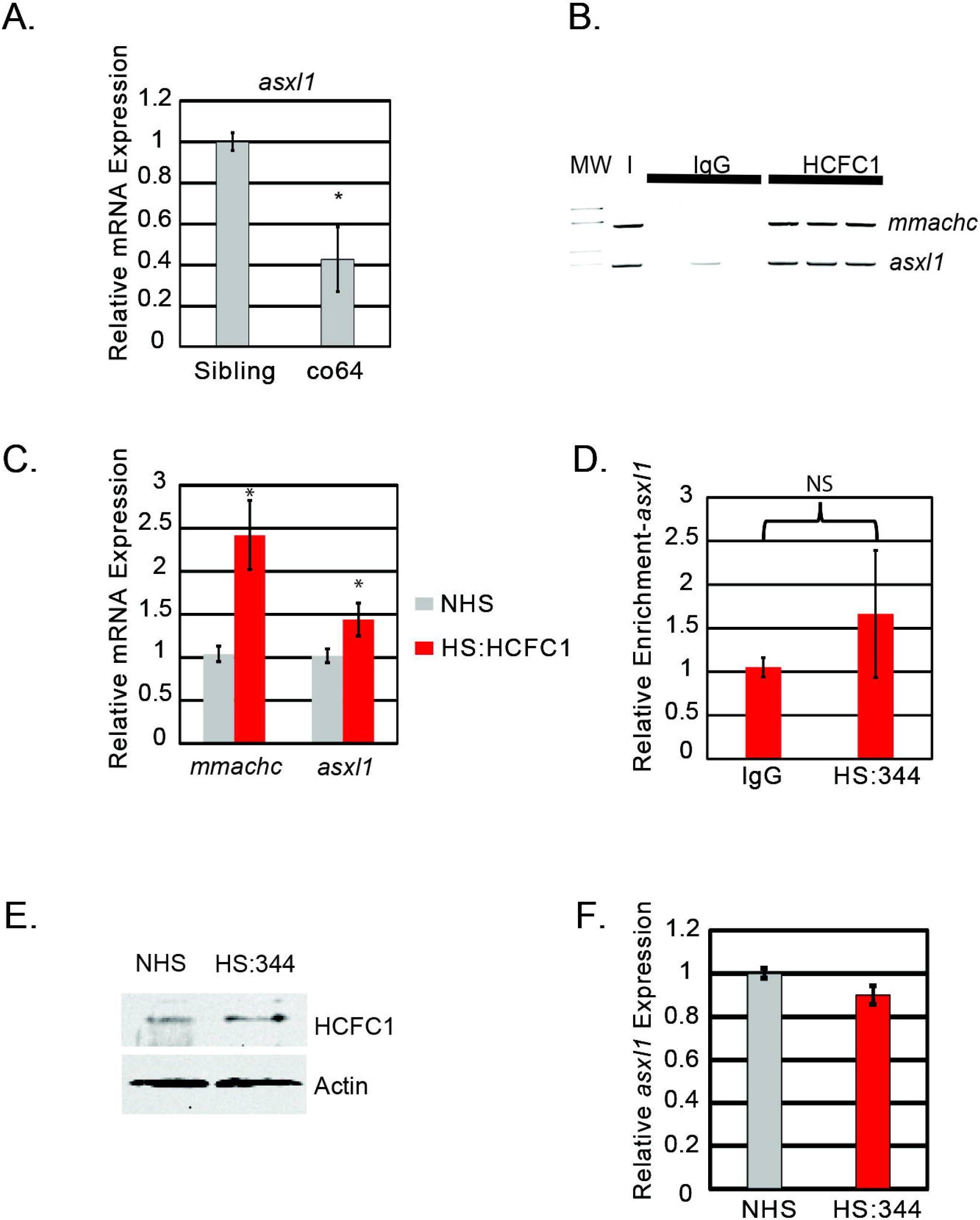
*cblX* mutations differentially regulate *asxl1* expression. A. Quantitative PCR (qPCR) was performed with total RNA isolated from whole brain homogenates of wildtype siblings or homozygous carriers of the *hcfc1a^co64^* allele (co64) to measure the expression level of *asxl1*. A total N=15 animals were used in two biological replicates. B. Chromatin Immunoprecipitation (ChIP) with semi-quantitative PCR was used to detect binding to the endogenous *mmachc* (+ control) or *asxl1* promoters. ChIP was performed using the *Tg*(*hsp701:HCFC1*). Binding was detected with anti-HCFC1 antibodies (HCFC1) relative to the IgG control (IgG). Lanes represent triplicate PCR from one biological replicate. Additional replicates were performed and validated using qPCR. Approximately 10% of the lysate (I) was utilized as an input control. N=20 heads per group were isolated and used for analysis. Assay was repeated on 3 independent occasions for a total N=60. C. qPCR was used to measure the expression of *mmachc* and *asxl1* in no heat shock (NHS) and heat shocked (HS:HCFC1) fish carrying the *Tg*(*hsp701:HCFC1*) transgene. Error bars represent standard error of the mean from biological replicates. Each biological replicate was performed with a pool of embryos from which total RNA was isolated. *p<0.05. N=20 heads per group across all replicates. D. ChIP was performed with anti-HCFC1 antibodies (HS:344) or IgG control (IgG) following a heat shock protocol with *Tg*(*hsp701:HCFC1 ^c.344C>T^)* larvae. N=20 heads per group were isolated and used for analysis for each biological replicate (3 replicates were performed). E. Western blot was performed with anti-HCFC1 antibodies or β-actin in no heat shock (NHS) or heat shocked *Tg*(*hsp701:HCFC1 ^c.344C>T^*) larvae (HS:344). N=4/group/biological replicate and performed in 2 biological replicates. F. qPCR analysis was utilized to determine the expression of *asxl1* in carriers of the *Tg*(*hsp701:HCFC1 ^c.344C>T^*) allele. *p<0.05. N=20 heads per group were isolated and used for analysis. Error bars represent standard error of the mean between biological replicates as indicated in materials and methods.

We next produced a heat shock transgene that would express the human *cblX* mutation (c.344C>T) variant (*Tg*(*hsp701:HCFC1^c.344C>T^*)) located in the kelch domain and previously described (Yu et al., 2013). Interestingly, this patient variant is the same variant produced in mice by Chern and colleagues. We performed ChIP using anti-IgG or anti-HCFC1 antibodies but did not detect binding to the *asxl1* promoter by qPCR (Fig 7D). Loss of enrichment was not the result of an inability of the antibody to bind the c.344C>T mutant variant, as western blot was used to validate the specificity of the HCFC1 antibody to the *cblX* variant protein (Fig 7E). Importantly, endogenous zebrafish Hcfc1 was detected in the no heat shock control as the antibody cross reacts with zebrafish Hcfc1 protein. Consistent with a loss of binding, we did not detect up-regulation of *asxl1* mRNA after heat shock protocol (Figure 7F). Collectively, these data suggest that HCFC1 binds to and activates the ASXL1 promoter, but that this function is disrupted in *cblX*.

### NPC/RGC number are independent of *asxl1* over-expression

Our data uncovered contrasting activity of Akt/mTor and differential *gfap* expression in the co60 and co64 alleles. These phenotypes correlate with unique signatures of *asxl1* expression, whereby high Akt/mTor activity is associated with increased *asxl1* expression. Furthermore, *cblX* mutations, including the c.344C>T variant, disrupt binding and activation to the ASXL1 promoter.

Therefore, we hypothesized that the mTor dependent phenotypes in the co60 allele were Asxl1 dependent. To test this, we injected *Asxl1* encoding mRNA into single cell embryos harboring either the Sox2 or Gfap EGFP reporter. We then monitored the number of total NPCs and RGCs by flow cytometry. As shown in Figure 8A, we did not detect any significant change in the number of Sox2+ or Gfap+ cells after over expression of *Asxl1*. We did not anticipate changes in the number of Sox2+ cells after Asxl1 injection because this is an overlapping phenotype present in the co60 and co64 alleles. Furthermore, we were not surprised that the number of Gfap+ cells was normal after injection because the co60 allele disrupts the expression of gfap and not cell number (Fig. 5E and 6C). However, we did not detect increased expression of *gfap* or *sox2* relative to non-injected control (Fig 8B). Importantly, injection of *Asxl1* encoding mRNA induced blood cell phenotypes consistent with a previously described function for this gene in hematopoiesis (Gjini et al., 2019; Uni and Kurokawa, 2018) (Supplemental Fig 1). The presence of blood cell phenotypes indicates functional translation of *Asxl1* mRNA after injection.

**Figure 8:**
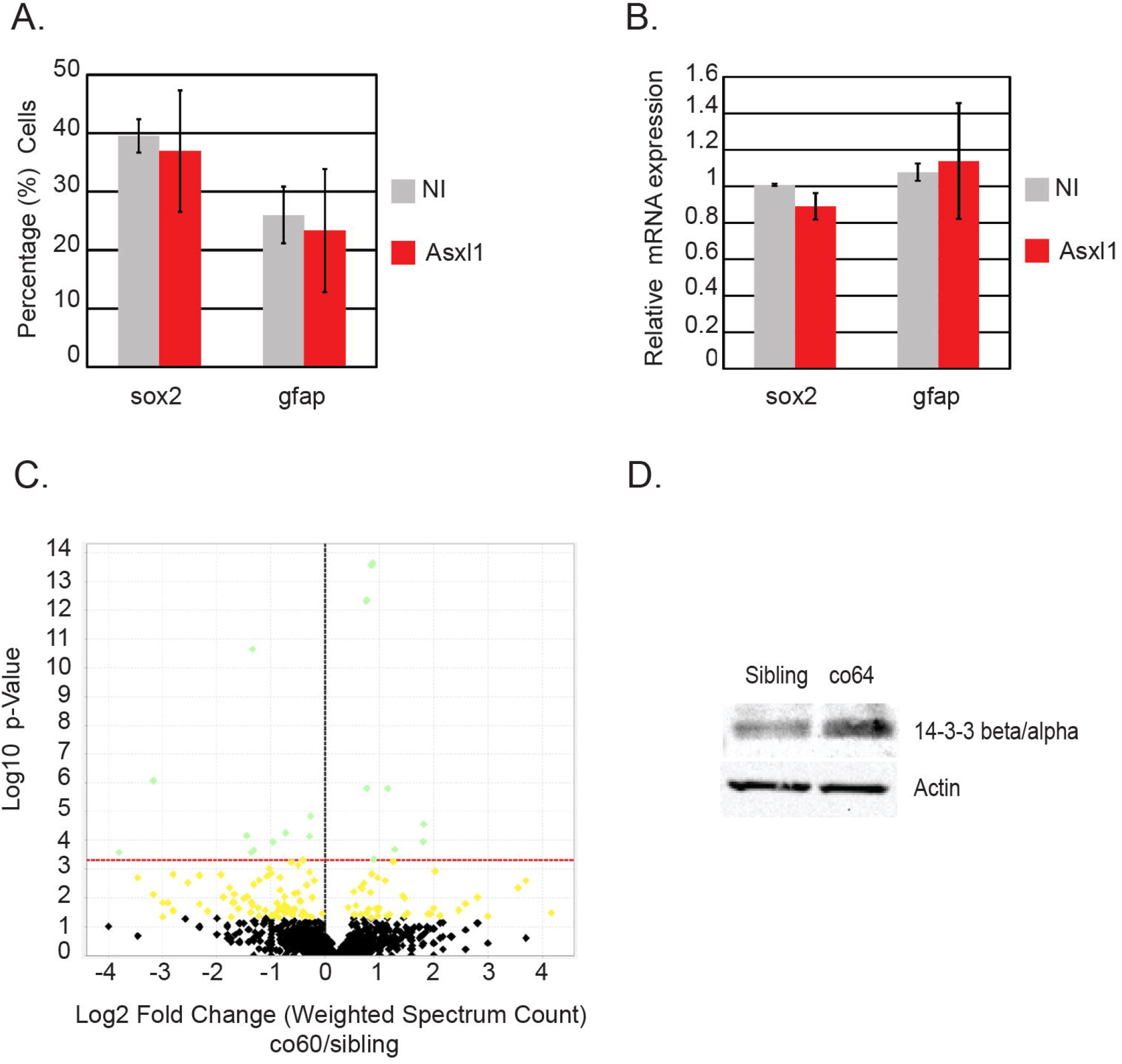
Forced expression of Asxl1 does not drive NPC/RGC development but proteomics reveals differential expression of 14-3-3 βα in each *cblX* allele. A. The number of Gfap+ or Sox2+ cells were quantified by flow cytometry in non-injected (NI), *Tg*(*gfap*:EGFP), or *Tg*(*sox2*:2A:EGFP) injected with mRNA encoding the murine *Asxl1* gene (Asxl1). Analysis was performed in two biological replicates. Error bars represent the standard error of the mean between biological replicates. N=14-20/group across 2 biological replicates. B. The relative expression of *gfap* or *sox2* was quantified in non-injected (NI) or wildtype embryos injected with mRNA encoding the murine *Asxl1* gene (Asxl1). N=12/group across 3 biological replicates. Error bars represent standard error of the mean from biological replicates. C. Mass spectrometry was performed to detect abnormal protein expression from N=13 brain homogenates obtained from sibling wildtype or carriers of the *hcfc1a^co60/+^*. Volcano plot describes the proteins that were significantly different between groups. D. Western blot analysis was performed to determine the expression of 14-3-3 βα in total brain homogenates of the co64 allele or sibling control. β-actin was used as a loading control. N=4/group/biological replicate. A total of 2 biological replicates were performed.

### Proteome analysis reveals abnormal expression of ribosomal proteins and 14-3-3βα in the co60 allele

Over-expression of *asxl1* did not alter RGC expression or number and therefore, we used proteomic analysis to identify novel mediators of Akt/mTor in the co60 allele. Our analysis identified a total of 2172 proteins that were differentially expressed with 159 of those proteins statistically different in the co60 allele. We identified many proteins belonging to biological processes such as hematopoiesis, erythrocyte development, the ribosome, and/or structural constituents of the ribosome. After secondary Benjamini-Hochberg correction, only 20 proteins were significantly different in the co60 allele relative to sibling control. Asxl1 protein was not detected as abnormal. Literature analysis of the 20 significant proteins demonstrated increased expression of ribosomal proteins: Rpl36a, Rpl38a, Rps7, Rpl18, Rpl9, Rpl30, Rpl18a, and Rpl3 (Supplementary Data File 1). Our identification of increased expression of proteins required for ribosome biogenesis is consistent with results obtained from a patient derived *cblX*-like knock-in allele (THAP11 gene), which adds additional face validity to our model system (Chern et al., 2022). Additional literature characterization of the remaining proteins revealed a statistically significant decrease in the expression of 14-3-3 βα (Fig. 8C). 14-3-3 proteins are a family of signaling proteins that bind to phosphorylated serine and threonine residues. They are known to regulate brain development (Cornell and Toyo-oka, 2017) and have been shown to reduce Akt phosphorylation at threonine 308 (Gómez-Suárez et al., 2016b). Based on the function of 14-3-3 proteins, we surmised that the co60 and co64 alleles would have opposite expression profiles as it relates to 14-3-3 βα. We used western blotting to determine the expression of 14-3-3 βα in the co64 allele and observed increased expression relative to sibling wildtype (Figure 8D), which contrasts with reduced expression in the co60 allele (Figure 8C). We next used the *Harmonizome* and The *Encyclopedia of DNA Elements* (ENCODE) databases to determine if HCFC1 binds to and regulates the expression of 14-3-3- β/α. 14-3-3- β/α is encoded by the *YWHAB* gene and we validated by database that HCFC1 binds to the *YWHAB* promoter (ENCODE Project Consortium, 2012; ENCODE Project Consortium, 2012; Luo et al., 2020; Rouillard et al., 2016). Thus, 14-3-3- β/α is an HCFC1 target gene with abnormal expression in the co60 and co64 alleles.

### The co60 allele accurately models the c.344C>T *cblX* variant

We observed two independent mTor signatures in the co60 and co64 alleles. One of which is associated with increased mTor and increased expression of proteins required for ribosome biogenesis. Thus, we questioned which of our alleles, and which mTor signature most closely resembles the molecular mechanisms underlying *cblX* syndrome. To begin to address this, we performed proteomics analysis using the *Tg*(*hsp701:HCFC1^c.344C>T^*) at 5 DPF. We observed a statistically significant up regulation of proteins essential for ribosome biogenesis and elongation initiation factors which are required for protein translation (Fig 9A). These data support and align with previous literature and therefore, provided additional face validity of heat shock HCFC1 transgenes to understand *cblX* syndrome. We plotted the average spectral count from 2 biological replicates for each of those proteins whose function is linked to ribosome biogenesis and translation. Each protein was increased in expression after induction of the c.344C>T patient variant (Figure 9A). We next hypothesized that an increase these proteins was a consequence of increased mTor signaling in the *Tg*(*hsp701:HCFC1^c.344C>T^*). We compared pS6 phosphorylation in heat shocked induced and non-heat shock control samples. We observed increased pS6 kinase and increased total S6, with equivalent levels of β-actin as a loading control (Fig 9B). We used β-actin as a control because we observed an increase in total S6 in multiple biological replicates. These data suggest that *cblX* mutations induce defects in the synthesis of proteins important for ribosomal biogenesis, consistent with previous literature.

**Figure 9:**
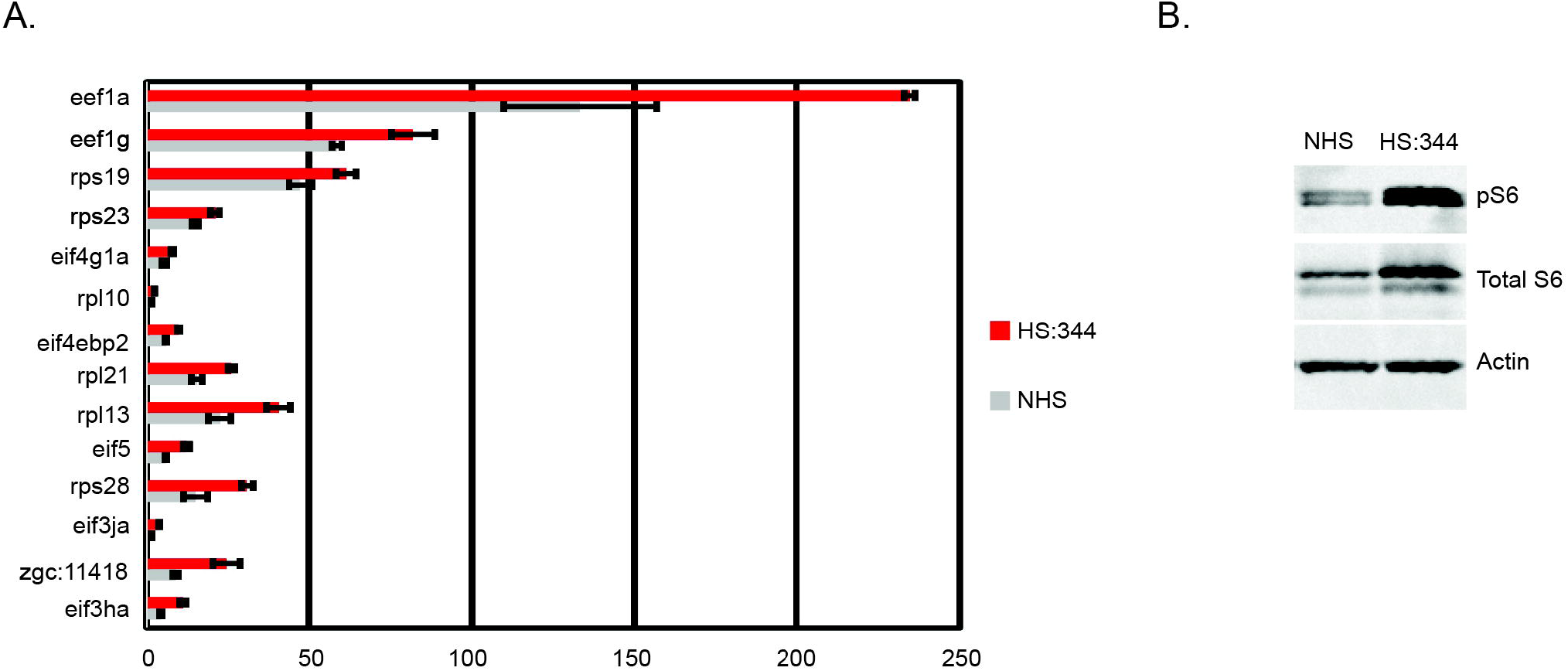
Proteomics demonstrates abnormal expression of ribosomal proteins and activation of Akt/mTor in the *Tg* (*hsp701:HCFC1^c.344C>T^*). A.Graph depicting average spectral counts of genes encoding proteins associated with ribosomal biogenesis and translation initiation. Proteomics was performed in biological triplicates and average spectral count in non-heat shock (NHS) and heat shocked (HS:344) animals carrying the *Tg*(*hsp701:HCFC1^c.344C>T^*) allele were averaged and plotted. Proteins shown were statistically up regulated in the c.344C>T allele. Error bars represent standard deviation between spectral counts obtained from biological replicates. A total of N=45 fish across 3 biological replicates were used for NHS and HS (HS:344) and N=46 fish across 3 biological replicates were used for NHS and HS *Tg*(*hsp701:HCFC1*) for analysis. B. Western blot was performed on whole brain homogenates with phospho-S6 kinase, total S6, or β-actin antibodies in no heat shock (NHS) or heat shocked animals carrying the *Tg*(*hsp701:HCFC1^c.344C>T^*) allele. Analysis was performed at 5 days post fertilization for both data in A and B. For western blot, a total N=4/group/biological replicate was utilized in 2 biological replicates.

## Discussion

Mutations in the *HCFC1* gene cause *cblX* syndrome and XLID (Gérard et al., 2015, 1; Huang et al., 2012; Jolly et al., 2015, 1; Koufaris et al., 2016b, 1; Scalais et al., 2017, 1; Yu et al., 2013). Various model systems have been developed and suggest that HCFC1 is critical for the function of NPCs (Castro et al., 2020; Chern et al., 2022; Huang et al., 2012; Jolly et al., 2015; Minocha and Herr, 2019; Minocha et al., 2016a; Minocha et al., 2016b; Quintana et al., 2017). Most recently, Chern and colleagues characterized the cellular and molecular phenotypes associated using a *cblX* patient knock-in allele (Chern et al., 2022). These analyses focused on craniofacial development and demonstrated only limited characterization of brain development, likely due to sub-viability. Despite limitations associated with viability, analysis of a related disorder caused by mutations in THAP11 (p.F80L), an HCFC1 interacting protein, revealed increased expression of proteins associated with ribosome biogenesis. These studies complement existing work in the zebrafish in which Castro and colleagued found brain phenotypes to be *mmachc* independent (Castro et al., 2020).

We sought to build on these findings by comparing the molecular and cellular phenotypes present in 2 zebrafish *hcfc1a* germline mutant alleles. We have previously characterized the cellular and molecular phenotypes present in a nonsense allele of *hcfc1a* (co60). However, *cblX* is the result of missense mutations and therefore, we created the co64 allele, the first homozygous viable germline mutant *in vivo*. We demonstrate that the co60 and co64 alleles differentially affect Hcfc1a protein expression. The co60 allele reduces expression of both N and C-terminal proteolytically cleavage fragments whereas the co64 allele does not, producing an N-terminal cleavage fragment with a missense mutation in the kelch domain. Multiple different mutations in HCFC1 have been reported in the literature including those that affect dosage and cause XLID (Huang et al., 2012). The co60 reduces protein expression and the dosage of HCFC1 resulting in cellular and molecular phenotypes that are unique from the co64 allele. Protein analysis confirmed the co64 allele did not affect dosage and genomic sequencing confirmed that the co64 allele resulted in a 2 amino acid modification in the N-terminal fragment. It is not surprising that mutations affecting dosage have distinct and overlapping phenotypes when compared with those that alter protein function rather than dosage. Particularly as it is related to HCFC1, as the protein is proteolytically cleaved and each fragment is known to have unique functions (Julien and Herr, 2003; Julien and Herr, 2004; Luciano and Wilson, 2002; Mangone et al., 2010). In addition, previous studies have established that unique mutations in other genes (*Gli3*) can cause different diseases (Johnston et al., 2005).

Our comparison of the co60 and co64 alleles revealed contrasting phenotypes at the level of Akt/mTor activation. Nonsense mutation of *hcfc1a*, which reduced expression of both N and C terminal proteolytic cleavage fragments, resulted in hyperphophorylation and activation of Akt and mTor. These data strongly support the observations from Chern and colleagues, as activation of mTor promotes translation and ribosome biogenesis, which was elevated in a knock-in allele of *THAP11* (*cblX*-like) (Chern et al., 2022). Thus, the co60 is a valid model system to study the mechanisms by which changes in protein dosage affect brain development and understand the mechanisms driving increased expression of ribosome biogenesis. Mutation of THAP11 causes a *cblX*-like syndrome (Quintana et al., 2017), but whether the underlying mechanisms associated with mutation of THAP11 are identical to *cblX* syndrome has not been elucidated. The expression level of proteins associated with ribosome biogenesis was not directly tested in *cblX*, but genetic complementation assays with germline mutants that cause ribosomopathies were performed. Here we more directly tested the role of ribosome biogenesis using the *Tg*(*hsp701:HCFC1^c.344C>T^*) transgene. This allele conditionally ubiquitously expresses the exact patient *cblX* mutation modeled by Chern and colleagues (Chern et al., 2022). We observed increased expression of proteins associated with ribosome biogenesis and translation indicating an underlying mechanism associated with the c.344C>T (p.A115V) patient mutation. Therefore, we provide face validity for the *Tg*(*hsp701:HCFC1^c.344C>T^*) as a model of *cblX* syndrome.

Interestingly, we observed hypophosphorylation of Akt/mTor after missense mutation of *hcfc1a*. These data contrast with the co60 allele and the *Tg*(*hsp701:HCFC1^c.344C>T^*) allele. We anticipate that the mechanisms present in the co64 allele also differ from the genetic knock-in created by Chern and colleagues (Chern et al., 2022). This is because the co64 deletes and in frame arginine and results in two amino acid changes within the kelch domain. Interestingly, despite differences in Akt/mTor signaling, the co60 and co64 have overlapping effects on the number of NPCs. These overlapping phenotypes provide evidence that Hcfc1a is essential for proper control of the NPC compartment and the presence of similar phenotypes across different alleles is indicative that the NPC phenotypes present in each allele are a direct result of mutation in Hcfc1a and not due to any unknown off-target effect. Knockdown of *hcfc1a* and *hcfc1b* by morpholino resulted in an increase in the number of NPCs (Quintana et al., 2017), providing additional validity of both the co60 and co64 alleles. These zebrafish studies are also supported by *in vitro* knockdown of Hcfc1, which resulted in an increase in NPCs (Jolly et al., 2015). Given the strong validity of each allele, our data raise the possibility that unique mutations in *hcfc1a* can lead to overlapping and individual phenotypes. We do not yet understand if individual disease variants will have differential regulation of mTor, but future studies are underway which may reveal unique signatures of Akt/mTor across individual disease variants. A role for mTor in *cblX* is further supported by a recent review that suggests the phenotypes of multiple neurodevelopmental disorders converge on the mTor pathway (Parenti et al., 2020).

Previously studies in the co60 allele uncovered increased expression of *asxl1*, which encodes a chromatin binding protein capable of interacting with Akt in the cytoplasm and promoting Akt phosphorylation (Castro et al., 2020; Youn et al., 2017). We previously used inhibitors of PI3K, an upstream regulator of Akt phosphorylation to indirectly link the NPC phenotypes present in the co60 allele to the Akt signaling pathway (Castro et al., 2020). Here we uncovered differential expression of *asxl1* in the co60 and co64 alleles. Most interestingly, was the fact that *asxl1* expression was correlated with hyper or hypo activation of Akt/mTor in each allele. We hypothesized that dysregulation of *asxl1* in *cblX* was the mechanism by which HCFC1 regulates RGC development. However, forced expression of *Asxl1* did not result in changes to the NPC or RGC populations. These data are inconsistent with morpholino mediated knockdown of *asxl1* in the co60 allele, which led to a full restoration of the number of NPCs (Castro et al., 2020). At the present time we cannot explain the mechanism by which morpholino knockdown of *asxl1* restored NPC phenotypes. But it is well-known that morpholinos can have off-target effects. Here we found that heat shock directed expression of wildtype HCFC1 caused an increase in *asxl1* expression, and we detected HCFC1 bound to the *asxl1* promoter. These data suggest that *asxl1* is a bonafide target of HCFC1. Since the *cblX* mutation p.Ala115Val disrupted binding of HCFC1 to this promoter, we investigated whether HCFC1 regulated NPC and RGC phenotypes via modulation of *asxl1* expression. However, forced expression of *Asxl1* did not affect the total numbers of NPCs or RGCs and had no effect on *gfap* expression. Interestingly, we also noted increased *asxl1* expression upon over expression of wildtype HCFC1. In our previous study we observed decreased *sox2* and *gfap* expression after over expression of wildtype HCFC1 (Castro et al., 2020). Thus, *asxl1* expression did not correlate with changes in *sox2* and *gfap* expression, which supports the data we observed after injection of *Asxl1* mRNA. We conclude that our collective data suggests that although *asxl1* expression is disrupted in cblX syndrome it is not likely to be the sole mediator of the Sox2+ NPC or Gfap+ RGC phenotypes in *cblX* or related disorders which include the co64 allele.

Our proteomics analysis of the co60 allele uncovered increased expression of proteins essential for ribosome biogenesis (Supplementary Data File 2). These data are consistent with proteomics analysis using the *Tg*(*hsp701:HCFC1^c.344C>T^*) allele where we identified increased expression of several elongation initiation factor proteins. Given that these two alleles demonstrate phenotypes consistent with the recent knock-in mouse models, we propose that both are valid systems to understand the mechanisms by which mutations in *cblX* cause disease. We also identified a second putative regulator of Akt signaling through proteomics, the 14-3-3- β/α protein. 14-3-3 proteins have been implicated in brain development (Cornell and Toyo-oka, 2017) and some members of the larger family have been shown to increase the number of NPCs (Cornell and Toyo-oka, 2017) and inhibit Akt phosphorylation (Gómez-Suárez et al., 2016b). We found reduced expression of 14-3-3- β/α in the co60 allele and increased expression in the co64 allele, which correspond with hyperactivated Akt/mTor and reduced Akt/mTor signaling, respectively. We validated through available data in the ENCODE database that HCFC1 binds to the *YWHAB* promoter. Future experiments to characterize the role of 14-3-3- β/α in *cblX* syndrome are warranted.

Collectively, our data suggest that distinct types of mutations, even in the same domain of HCFC1, can have contrasting phenotypes at the level of Akt/mTor, *asxl1* expression, and *gfap* expression. Initially, the presence of contrasting phenotypes may call into question the physiological relevance of the co64 allele because the phenotypes in the co60 allele, such as increased synthesis of proteins required for ribosomal biogenesis, have been validated with recent patient derived models (Chern et al., 2022). However, cell type specific deletion of *Hcfc1* in the Nkx+2.1+ sub-compartment of NPCs reduces Gfap expression (Minocha and Herr, 2019), data that supports reduced Gfap expression in the co64 allele. Interestingly, the THAP11 p.F80L allele demonstrated decreased Gfap expression after knock-in providing additional evidence for a role of HCFC1 and its partner THAP11 in the regulation of GFAP expression. The F80L mutation also has increased expression of proteins needed for ribosomal biogenesis. Interestingly, the mTor dependent regulation of *gfap* expression in the co60 allele combined with decreased mTor and *gfap* expression in the co64 allele further substantiates the validity for the co64 allele as a model to inform about *cblX* syndrome. Unfortunately, the co64 could not be directly rescued because over expression of HCFC1 and heat shock expression of HCFC1 cause independent phenotypes (Castro et al., 2020; Quintana et al., 2017) complicating the interpretation of restoration assays. However, the co60 and co64 alleles have been outcrossed for 15-20 generations each, limiting the potential effects of off-target effects.

In conclusion, our study uncovers differential regulation of Akt/mTor across different types of germline mutations in the zebrafish *hcfc1a* gene. Our data correlate contrasting levels of Akt/mTor to unique cellular defects of the Gfap+ RGC cellular compartment. Most intriguingly, our work suggests that Asxl1 and 14-3-3βα may converge to regulate Akt activity in an HCFC1 deficient background. Collectively, this work provides a fundamental mechanism (Akt/mTor) by which increased synthesis of proteins required for ribosome biogenesis occurs in *cblX* syndrome. However, our work also suggests that independent mutations with unique effects on overall HCFC1 protein expression/function could cause disease by unique mechanisms that may or may not include Akt/mTor perturbation. Future studies characterizing the function of unique disease variants in various animal models are underway.

## Material and Methods

### Experimental model and subject details

For the experiments described, embryos were produced from natural spawnings of the following lines: *hcfc1a^co64^*, *hcfc1a^co60/+^*, *Tg*(*sox2:2A:EGFP*), *Tg*(*gfap:EGFP*), *Tg*(*hsp701:HCFC1*), *Tg*(*hsp701:^HCFC1c.344C>T^*), AB, or Tupfel long fin. Embryos were maintained in E3 media as described by the protocols on the Zebrafish Information Network (ZFIN) at 28° with a 14/10 light:dark cycle. All procedures were approved by the Institutional Animal Care and Use Committee at the University of Texas El Paso, protocol number 811869-5. Methods for euthanasia and anesthesia were performed according to guidelines from the 2020 American Veterinary Medical Association guidebook.

The *hcfc1a^co64^* allele was created as previously described (Castro et al., 2020) with an identical guide RNA and with equivalent CRISPR/Cas9 editing technology as described. The *hcfc1a^co64^* was created at the same time as the afore published *hcfc1a^co60/+^* allele except that the insertion/deletion created, produced a second allele with the sequences described. The offspring of the *hcfc1a*^co64^ allele were generated from a single founder (F_0_), which was outcrossed with 3 independent wildtype fish to generate a minimum of 3 families of F1 carriers. Each family consisted of approximately 20 total fish with equal numbers of males and females. To generate subsequent generations, we outcrossed a minimum of 3-F1 individuals with wildtype to obtain a minimum of 3 families of F2 carriers. We subsequently outcrossed F2 carriers (minimum of 3) with wildtype (AB) fish to produce an F3 generation of approximately 3 total families with equal numbers of males and females. Sanger sequencing confirmed mutation of the allele and experiments were initiated in the F3 generation. The *Tg*(*hsp701:HCFC1^c.344C>T^*) was created using Gateway cloning technology as previously described (Castro et al., 2020; Kwan et al., 2007). Vectors utilized for the HCFC1 open reading frame were previously described (Quintana et al., 2014; Quintana et al., 2017). The experiments described herein were performed in the F2 generation, which was produced from a single founder (F_0_). The positive F_0_ carrier was outcrossed with wildtype (AB) to produce 2 families of F1 individuals and a minimum of 3 carriers of the F1 generation were outcrossed to produce 3 families of F2 carriers that were utilized for the experiments described. All shared resources will be shared via ZFIN.

### Genotyping of indicated zebrafish lines

Genotyping was performed with DNA from excised larval tissue or fin clips (adults). Tissue was lysed in 50mM sodium hydroxide (Fisher Scientific) for 15 minutes at 95° Celsius and pH adjusted with 1M Tris-HCl. Primers pairs were developed for each allele that specifically bind to and amplify the mutated allele and will not amplify the wildtype allele (Castro et al., 2020). For the *hcfc1a*^co60/+^ the primers specific to the mutated allele are FWD: CCAGTTCGCCTTTTTGTTGT and REV: ACGGGTGGTATGAACCACTGGC. PCR annealing was performed at 64°. For the *hcfc1a*^co64^ allele, the mutant allele was amplified by standard PCR at an annealing temperature of 64° with FWD: CCAGTTCGCCTTTTTGTTGT and REV: CTGGAGGGATGTTCATAACGG). For identification of the wildtype allele, the following primers were utilized: FWD: CCAGTTCGCCTTTTTGTTGT and REV: TCCCCACGAACGGCTGGTAT.

Genotyping of the *Tg*(*hsp701*:*HCFC1*) allele was performed with the following primers: FWD: TGAAACAATTGCACCATAAATTG present in the *hsp701* promoter and REV: CGTCACACACGAAGCCATAG in the *HCFC1* open reading frame. PCR for HCFC1 genotyping was performed at 60° annealing temperature with standard extension times. The two heat shock alleles were genotyped with overlapping primer pairs. Genotyping and validation of this allele was previously described (Castro et al., 2020)

### Protein Isolation and western blotting

Protein was isolated from the excised brains of larvae at 5 DPF. Brains were homogenized using a pestle in 1X Radio-Immuno-Precipitation-Assay (RIPA) (Fisher Scientific) or 1X Cell Lysis Buffer (Cell signaling). Protease inhibitors were added at a final 1X concentration (Fisher Scientific). Supernatant was collected after a 10-minute centrifugation at 10,000XG in a cold centrifuge. Protein concentration was quantified using Precision Red (Cytoskeleton) according to manufacturer’s instructions. Western blots were performed using standard techniques using the Novex Tris-Glycine-SDS system (Fisher Scientific). Protein loading was quantified using Ponceau S (Fisher Scientific) and membranes were blocked with 5% ECL Prime Blocking Agent (Fisher Scientific). Primary antibody concentrations are as follows: *pAkt* (Thr308) (Cell Signaling Cat. # D25E6) (anti-rabbit) 1:3000, *Akt* (pan) (Cell Signaling Cat. # C67E7) (anti-rabbit) 1:3000, 14-3-3-beta/alpha (Cell Signaling Cat # 9636S) (anti-rabbit) 1:3000, S6 Ribosomal (5G10) (Cell Signaling Cat # 2217S) (anti-rabbit) 1:3000, P-S6 Ribosomal Protein (S235/236) (Cell Signaling Cat # 4858S) (anti-rabbit) 1:3000, Asxl1 (Cell Signaling Cat # 52519S) (anti-rabbit) 1:3000, Beta-Actin (Sigma/Millipore Cat # A5441 clone AC-15) (anti-mouse) 1:5000, N-terminal HCFC1 (Sigma/Millipore Cat # QC10816) (anti-rabbit) 1:500, C-terminal HCFC1 (Bethyl labs Cat # 50- 156-0286 purchased through Fisher Scientific) (anti-rabbit) 1:100. Secondary antibodies were utilized at a concentration of 1:20,000 and chemiluminescence was performed with ECL Primer Western Blotting detection reagent (Fisher Scientific). Imaging of membranes was performed using an iBright chemiblot system.

### RNA isolation and quantitative real time PCR

RNA was isolated from embryos at the indicated time points using Trizol (Fisher Scientific) according to manufacturer’s protocol. Reverse transcription was performed using Verso cDNA synthesis (Fisher Scientific) and total RNA was normalized across all samples. For each biological replicate (2-5/experiment) PCR was performed in technical triplicates using an Applied Biosystem’s StepOne Plus machine with Applied Biosystem’s software. Analysis was performed using 2^ΔΔct^. Sybr green (Fisher Scientific) based primer pairs for each gene analyzed are as follows: *hcfc1a fwd*: ACAGGGCCTAACACAGGTTG, *hcfc1a rev*: TCCTGTGACTGTGCCAAGAG, *hcfc1b fwd*: GGATGGGTCCCTCTGGTTAT, *hcfc1b rev*: CGGTAACCATCTCGTCCACT, *mmachc fwd*: GCTTCGAGGTTTACCCCTTC, *mmachc rev*: AGGCCAGGGTAGGGTCCTG, *asxl1 fwd*: CCAGAGCTGGAAAGAACGTC, *asxl1 rev*: ACATCTCCAGCTTCGCTCAT, *rpl13a* fwd: TCCCAGCTGCTCTCAAGATT, *rpl13a* rev: TTCTTGGAATAGCGCAGCTT.

### Proteomics sample preparation

Samples were isolated from whole brain as described above and proteins were precipitated from lysis buffer using Trichloroacetic acid (TCA) as follows: 50μl of 100% TCA (Sigma/Millipore – cat# T6399-5G) was added to 200 μl of protein sample at a final concentration of 20%, then incubated at 4°C for 10 minutes. The mixture was centrifuged at 14,000 RPM for 5 minutes. Pellet was resuspended in 200μl of 100% LC/MS grade Acetone and centrifuged at 14,000 RPM for 5 minutes. Supernatant was discarded. This process was repeated two more times to ensure removal of residual detergents. After the third wash, the pellet was heated on a heating block at 95°C for 2 minutes to ensure acetone evaporation. Samples were stored at −80°C and resuspended in 8M Urea prior enzymatic digestion with trypsin.

Enzymatic tryptic digestion of proteins was obtained by using iST Sample preparation kit (Preomics, cat # iST 96x 00027). In summary, 10μl of Lysis buffer was added per 1μg of protein from zebrafish larvae brains and placed in a heating block (80°C; 20min, mixing every 5 min) for protein denaturing, alkylation, and reducing disulfide bonds within it. Droplets were the subject of a short spin (room temperature; 300rcf; 10 sec). Samples were sonicated using 10 cycles: at 30 seconds on/off. After sonication of the samples, 50 μL of resuspended of enzyme was added to samples for protein digestion. Microtubes containing samples were gently vortexed and placed in a heating block at 37°C for 90 minutes. Samples were gently mixed, every 10 min, during the 90- minute incubation period. 100μl of stop solution were added to samples before shaking and mixing the entire solution by pipetting up and down at least 10 times. Adapters and cartridges were placed on top of 1.5mL waste microtubes and samples were transferred to the cartridge. Cartridge containing samples were centrifuged at 3,800 rcf for 1 minute and samples were washed once with 200μL of Wash 1 and Wash 2 solutions. Samples were centrifuged at 3,800 rcf for 1 minute between wash steps. Centrifuged cartridges containing protein samples were transferred to a new collection tube prior to elution with two cycles of 100μL of Elution solution obtained by centrifugation at 3,800 rcf; 1 minute each cycle. Flow-through, containing peptides was placed in a vacuum evaporator at 45°C, 100 mTorr, for 90 min (Savant; Thermo Fisher Scientific), until completely dry. Eluted peptides were stored at −80°C until ready for analysis.

### Liquid chromatography–tandem mass spectrometry (LC/MS)

Resultant complex peptide mixtures were resuspended with 4% acetonitrile (ACN), 0.1% formic acid (FA), both LC/MS grade reagents at 1 μg/μL. One microliter of resuspended samples was loaded onto an Acclaim PepMap rapid separation LC column (75μm x 50cm nanoViper, PN 164942, Thermo Scientific), which was previously equilibrated with 4% solvent B (100% acetonitrile, 0.1% formic acid), and 96% solvent A (100% H_2_O, 0.1% formic acid). The multi-step gradient was started with peptides being loaded at a flow rate of 0.5μL/min for 15 minutes using a Dionex Ultimate 3000 RSLCnano (Thermo Scientific). The flow rate was decreased to 0.3μL/min over 15 min, and the solvent B decreased to 20% at 115 minutes, then increase to 32% at 135 minutes, and finally increased to 95% solvent B over 1 minute. To remove any additional remaining peptides, 95% solvent B was held for 4 minutes at a flow rate of 0.4μL/min. The flow rate was increased to 0.5μL/min over 1 minute and decreased to 4% solvent B. The column was re-equilibrated at 4% solvent B for 39 minutes and maintained at a flow rate 0.5μL/min for a total of 180 minutes total run time. The column was kept at flat 55°C during the entire analysis. One blank injection was performed in between biological samples using a 60 min seesaw wash using 4-95% ACN gradients.

Peptide data was acquired with a Q-Exactive Plus Hybrid Quadrupole-Orbitrap Mass Spectrometer (Thermo Scientific) in positive mode, set to data dependent MS^2^. Full MS ions were collected at resolution of 70,000, AGC target of 3e^6^, with a scan range of 375 to 1500 *m/z*. Ions were fragmented with NCE set to 27 and collected at a resolution of 17,500 with AGC target at 1e^5^, 2 *m/z* isolation window, maximum IT set to 60 ms, loop count 10, TopN 10. Charged exclusion ions were set to unassigned, 1, 6 – 8, >8.

### Bioinformatics data analysis

After LC/MS analysis, Proteome Discover (PD) 2.5.0.400 (Fisher Scientific) was utilized to identify the proteins from each peptide mixture. The database for Danio rerio was downloaded from UniProtKB; http://www.uniprot.org/ on 21 October 2021 with a database 61,623 sequences. A contaminant dataset was run in parallel composed of trypsin autolysis fragments, keratins, standards found in CRAPome repository1 and in-house contaminants. PD analysis parameters are as follows: false-discovery rate (FDR) of 1%, HCD MS/MS, fully tryptic peptides only, up to 2 missed cleavages, parent-ion mass of 10 ppm (monoisotopic); fragment mass tolerance of 0.6 Da (in Sequest) and 0.02 Da (in PD 2.1.1.21) (monoisotopic). Two-high confidence peptides per protein were applied for identifications. PD dataset was processed through Scaffold Q+S 5.0.1. Scaffold (Proteome Software, Inc., Portland, OR 97219, USA) was used to probabilistically validate protein identifications derived from MS/MS sequencing results using the X!Tandem2 and Protein Prophet computer algorithms3. Data was transferred to Scaffold LFQ (Proteome Software, Portland, Oregon, USA) which was used to validate and statistically compare protein identifications derived from MS/MS search results. A protein threshold of 95%, peptide threshold of 95%, and a minimum number of 2 peptides were used for protein validation. Normalized weighted spectral counts were used when comparing the samples. To ascertain p-values, Fisher’s Exact was run with a control FDR level q * .05 with standard Benjamini-Hochberg correction.

### Flow cytometry

To identify mutants, we adapted a protocol whereby tissue was excised from caudal most tip of the tail from larvae at 2-3 DPF (Kosuta et al., 2018). Tissue was collected on a filter paper and genotyped as described in the sections above. Equal numbers of mutants and wild type larvae were euthanized according to standard procedures, then heads were excised from tails to create a whole brain homogenate. Dissociation of heads was followed exactly according to previous protocol (Bresciani et al., 2018). Briefly, tissue was dissociated using Trypsin-EDTA (0.25%) with collagenase (100mg/mL), applying heating at 30° Celsius and disturbing tissue by pipetting up and down. The reaction was stopped by adding lamb serum (Fisher Scientific) and cells were harvested through centrifugation (5 minutes at 700g at room temperature). The cells were resuspended in 1X phosphate buffered saline (Fisher Scientific) and filtered through 40-70uM filter before performing flow cytometry. Analysis was performed with Kaluza software at a fixed rate of 10,000 cell events for a maximum of 3 minutes.

### Immunohistochemistry

Larvae were fixed at the indicated time points in 4% paraformaldehyde (Electron Microscopy Sciences) for minimum of 1 hour at room temperature. Genotyping was performed as described (Kosuta et al., 2018). Brain tissue was embedded in 1.5% agarose (Fisher Scientific) produced in 5% sucrose (Fisher Scientific). Embedded blocks were incubated overnight in 30% sucrose (Fisher Scientific) and then snap frozen with dry ice before cryosectioning (20-30μM). Validated transgenic reporter animals were utilized for visualization of Sox2+ and Gfap+ cells (ZDB-TGCONSTRCT-070410-2). For DNA content staining, Hoescht (2ug/ml) was utilized. Hoescht stain was performed according to manufacturer’s protocol available as part of the EdU Click-It technology kit (Fisher Scientific). For pulse labeling, larvae were incubated in 20 mM 5-ethynyl-2′-deoxyuridine (EdU) (Fisher Scientific) diluted in 10% dimethyl sulfoxide (DMSO) (Fisher Scientific) for 30 min at room temperature (RT) prior to fixation. EdU was visualized using the EdU Click-It technology. All slides were cover slipped using Vectashield (Vector Laboratories) and imaged on a Zeiss LSM 700 at 20×–63× magnification.

### Chromatin Immunoprecipitation (ChIP)

ChIP was performed as previously described (Quintana et al., 2011) except that larvae were disassociated into a single cell homogenate prior to cell fixation with formaldehyde. Briefly, equal numbers of non-heat shocked (NHS) and heat-shocked (HS) larvae in either line, *Tg*(*hsp701*:*HCFC1*) or *Tg(hsp701:HCFC1 ^c.344C>T^*), were euthanized according to standard procedures, then heads were excised from tails to create a whole brain homogenate. Tissue was dissociated as described above for flow cytometry. Shearing was performed in a 200 μl volume of 50 mM Tris pH 8.0 (Millipore/Sigma) by adding 40 units of Micrococcal nuclease (Fisher Scientific) for 10 minutes at 37°C. Ethylenediaminetetraacetic acid (EDTA-Fisher Scientific) (10 mM) was used to stop the reaction and nuclei were lysed in 1% sodium dodecyl sulfate (SDS-Fisher Scientific). ChIP was performed with anti-HCFC1 antibodies (Millipore Sigma AV38600), Anti-IgG (Santa Cruz Biotechnology, sc-2025). The collected precipitates were washed and de-crosslinked and the DNA was purified using standard protocols (Quintana et al., 2011). qPCR was used to determine enrichment using primers designed to the 1 kb region upstream of each start site/gene of interest. Primers to *GAPDH* putative promoter were used as a negative control and each set of enrichment was normalized to GAPDH PCR or input as described (Quintana et al., 2011). Primers used for SYBR green (Fisher Scientific) based PCR are as follows: *Ywhaba* fwd: AAAGCGTCTCGGTGTGTAGG, *Ywhaba* rev: GGTAGGATGCCAAAATCGAA, *Ywhabb* fwd: TTGCTCTCCGGTAGTTCTGG, *Ywhabb* rev: TTCGTTTCCCATGCTTCTCT, *Gapdh* fwd: GCCTGATTTGGTTGTGTCCT, *Gapdh* rev: CCAATTGGGAAATGCTTGAG, *Asxl1* fwd: GGCTGTAGGAGCGACTGAAG, *Asxl1* rev: TAAACACACACAGGGCGAAG.

### Heat shock and drug treatment assays

Heat shock was performed as previously described (Castro et al., 2020; Hudish et al., 2013). Briefly, heat shock was initiated at 24 hours post fertilization (HPF) and performed twice daily every 8-12 hours until 5 DPF. Heat shock was performed for a duration of 30-45 minutes by incubating dishes at 38°. For rapamycin treatment (Selleck Chemicals), the drug was dissolved in 100% DMSO (Fisher Scientific) and embryos were treated at 24 HPF and 72 HPF with a 0.8 uM concentration for a period of 24 hours. Concentration was determined empirically using a gradient coupled with gene expression of *ccne1*. The concentration utilized reduced *ccne1* expression, without any effect on larval viability. Media was removed and no treatment was applied at 48 and 96 HPF. Total treatment time during the 5-day period is 48 hours. Total time with treatment off is 72 hours.

## Acknowledgements

The authors would like to thank Drs. Bruce Appel and Tamim Shaikh for their sharing of model organisms associated with this manuscript. Special thanks to Yahir Davila, Jennifer Davila, Isaiah Perez, and Nayeli Reyes-Nava for their role in genotyping and animal husbandry. Proteomics analysis was performed in conjunction with the UTEP core facilities, and we graciously thank the directorship of Dr. Igor Almeida and Dr. Renato Aguilera. Additional support was provided by the College of Science and Drs. Robert Kirken and Michael Kenney. Additional feedback and commentary were provided by Dr. Charlotte Vines prior to submission. All members of the Quintana laboratory from 2019-present aided in animal husbandry and fish care to help facilitate the work described.

## Data availability

All data associated with this publication will be publicly available through database or contact with the corresponding author upon publication

## Author Contributions

VLC helped conceive and design the goals and experiments of the paper, performed flow cytometry in the co60 allele, conceived and executed the rapamycin inhibition assays, helped optimize protein isolation and performed proteomics in the co60 allele, performed injections and analysis with *Asxl1* encoding mRNA and transgenic alleles, performed immunohistochemistry and RNA expression analysis associated with restoration experiments, performed all ChIP assays, and wrote portions of the methods and discussion sections with edits to introduction and results. DP optimized all antibodies used in combination with AMQ and performed all western blots associated with Akt/mTor and HCFC1, performed analysis of NPC and RGC number/expression in the co64 allele, qPCR analysis of *hcfc1a*, *hcfc1b*, and *asxl1* in the co64, and optimized aspects of allele specific PCR used for genotyping. VV assisted DP in the cellular analysis and flow cytometry. ILE, BIA, and CCE performed proteomics and analysis. AMQ performed western blots, genotyped embryos for proteomics, evaluated data for flow cytometry, optimized gating for flow cytometry, cloned and prepared Asxl1 mRNA, designed experiments to optimize rapamycin treatment using empirical data, performed qPCR associated with gene expression, performed data analysis and interpretation of all figures, developed the heat shock transgenes, wrote the manuscript (introduction and results with edits to discussion and material and methods), made the figures, and performed western blots to validate heat shock transgene as shown.

## References

An, S., Park, U.-H., Moon, S., Kang, M., Youn, H., Hwang, J.-T., Kim, E.-J. and Um, S.-J. (2019). Asxl1 ablation in mouse embryonic stem cells impairs neural differentiation without affecting self-renewal. Biochem. Biophys. Res. Commun. 508, 907–913.

Bresciani, E., Broadbridge, E. and Liu, P. P. (2018). An efficient dissociation protocol for generation of single cell suspension from zebrafish embryos and larvae. MethodsX 5, 1287–1290.

Castro, V. L., Reyes, J. F., Reyes-Nava, N. G., Paz, D. and Quintana, A. M. (2020). Hcfc1a regulates neural precursor proliferation and asxl1 expression in the developing brain. BMC Neurosci 21, 27.

Chern, T., Achilleos, A., Tong, X., Hill, M. C., Saltzman, A. B., Reineke, L. C., Chaudhury, A., Dasgupta, S. K., Redhead, Y., Watkins, D., et al. (2022). Mutations in Hcfc1 and Ronin result in an inborn error of cobalamin metabolism and ribosomopathy. Nat Commun 13, 134.

Cornell, B. and Toyo-oka, K. (2017). 14-3-3 Proteins in Brain Development: Neurogenesis, Neuronal Migration and Neuromorphogenesis. Front Mol Neurosci 10, 318.

Dehaene, H., Praz, V., Lhôte, P., Lopes, M. and Herr, W. (2020). THAP11F80L cobalamin disorder-associated mutation reveals normal and pathogenic THAP11 functions in gene expression and cell proliferation. PLoS One 15, e0224646.

Dejosez, M., Krumenacker, J. S., Zitur, L. J., Passeri, M., Chu, L.-F., Songyang, Z., Thomson, J. A. and Zwaka, T. P. (2008). Ronin is essential for embryogenesis and the pluripotency of mouse embryonic stem cells. Cell 133, 1162–1174.

Dejosez, M., Levine, S. S., Frampton, G. M., Whyte, W. A., Stratton, S. A., Barton, M. C., Gunaratne, P. H., Young, R. A. and Zwaka, T. P. (2010). Ronin/Hcf-1 binds to a hyperconserved enhancer element and regulates genes involved in the growth of embryonic stem cells. Genes Dev. 24, 1479–1484.

ENCODE Project Consortium (2012). An integrated encyclopedia of DNA elements in the human genome. Nature 489, 57–74.

Gabut, M., Bourdelais, F. and Durand, S. (2020). Ribosome and Translational Control in Stem Cells. Cells 9, 497.

Gérard, M., Morin, G., Bourillon, A., Colson, C., Mathieu, S., Rabier, D., Billette de Villemeur, T., Ogier de Baulny, H. and Benoist, J. F. (2015). Multiple congenital anomalies in two boys with mutation in HCFC1 and cobalamin disorder. Eur J Med Genet 58, 148–153.

Gjini, E., Jing, C.-B., Nguyen, A. T., Reyon, D., Gans, E., Kesarsing, M., Peterson, J., Pozdnyakova, O., Rodig, S. J., Mansour, M. R., et al. (2019). Disruption of asxl1 results in myeloproliferative neoplasms in zebrafish. Dis Model Mech 12, dmm035790.

Gómez-Suárez, M., Gutiérrez-Martínez, I. Z., Hernández-Trejo, J. A., Hernández-Ruiz, M., Suárez-Pérez, D., Candelario, A., Kamekura, R., Medina-Contreras, O., Schnoor, M., Ortiz-Navarrete, V., et al. (2016a). 14-3-3 Proteins regulate Akt Thr308 phosphorylation in intestinal epithelial cells. Cell Death Differ 23, 1060–1072.

Gómez-Suárez, M., Gutiérrez-Martínez, I. Z., Hernández-Trejo, J. A., Hernández-Ruiz, M., Suárez-Pérez, D., Candelario, A., Kamekura, R., Medina-Contreras, O., Schnoor, M., Ortiz-Navarrete, V., et al. (2016b). 14-3-3 Proteins regulate Akt Thr308 phosphorylation in intestinal epithelial cells. Cell Death Differ 23, 1060–1072.

Huang, L., Jolly, L. A., Willis-Owen, S., Gardner, A., Kumar, R., Douglas, E., Shoubridge, C., Wieczorek, D., Tzschach, A., Cohen, M., et al. (2012). A Noncoding, Regulatory Mutation Implicates HCFC1 in Nonsyndromic Intellectual Disability. Am. J. Hum. Genet. 91, 694–702.

Hudish, L. I., Blasky, A. J. and Appel, B. (2013). miR-219 regulates neural precursor differentiation by direct inhibition of apical par polarity proteins. Dev. Cell 27, 387–398.

Inoue, D., Fujino, T., Sheridan, P., Zhang, Y.-Z., Nagase, R., Horikawa, S., Li, Z., Matsui, H., Kanai, A., Saika, M., et al. (2018). A novel ASXL1-OGT axis plays roles in H3K4 methylation and tumor suppression in myeloid malignancies. Leukemia 32, 1327–1337.

Johnston, J. J., Olivos-Glander, I., Killoran, C., Elson, E., Turner, J. T., Peters, K. F., Abbott, M. H., Aughton, D. J., Aylsworth, A. S., Bamshad, M. J., et al. (2005). Molecular and Clinical Analyses of Greig Cephalopolysyndactyly and Pallister-Hall Syndromes: Robust Phenotype Prediction from the Type and Position of GLI3 Mutations. Am J Hum Genet 76, 609–622.

Jolly, L. A., Nguyen, L. S., Domingo, D., Sun, Y., Barry, S., Hancarova, M., Plevova, P., Vlckova, M., Havlovicova, M., Kalscheuer, V. M., et al. (2015). HCFC1 loss-of-function mutations disrupt neuronal and neural progenitor cells of the developing brain. Hum. Mol. Genet. 24, 3335–3347.

Julien, E. and Herr, W. (2003). Proteolytic processing is necessary to separate and ensure proper cell growth and cytokinesis functions of HCF-1. EMBO J. 22, 2360–2369.

Julien, E. and Herr, W. (2004). A switch in mitotic histone H4 lysine 20 methylation status is linked to M phase defects upon loss of HCF-1. Mol. Cell 14, 713–725.

Kapuria, V., Röhrig, U. F., Bhuiyan, T., Borodkin, V. S., van Aalten, D. M. F., Zoete, V. and Herr, W. (2016). Proteolysis of HCF-1 by Ser/Thr glycosylation-incompetent O-GlcNAc transferase:UDP-GlcNAc complexes. Genes Dev 30, 960–972.

Kosuta, C., Daniel, K., Johnstone, D. L., Mongeon, K., Ban, K., LeBlanc, S., MacLeod, S., Et-Tahiry, K., Ekker, M., MacKenzie, A., et al. (2018). High-throughput DNA Extraction and Genotyping of 3dpf Zebrafish Larvae by Fin Clipping. J Vis Exp.

Koufaris, C., Alexandrou, A., Tanteles, G. A., Anastasiadou, V. and Sismani, C. (2016a). A novel HCFC1 variant in male siblings with intellectual disability and microcephaly in the absence of cobalamin disorder. Biomed Rep 4, 215–218.

Koufaris, C., Alexandrou, A., Tanteles, G. A., Anastasiadou, V. and Sismani, C. (2016b). A novel HCFC1 variant in male siblings with intellectual disability and microcephaly in the absence of cobalamin disorder. Biomedical Reports 4, 215–218.

Kwan, K. M., Fujimoto, E., Grabher, C., Mangum, B. D., Hardy, M. E., Campbell, D. S., Parant, J. M., Yost, H. J., Kanki, J. P. and Chien, C.-B. (2007). The Tol2kit: a multisite gateway-based construction kit for Tol2 transposon transgenesis constructs. Dev. Dyn. 236, 3088–3099.

Luciano, R. L. and Wilson, A. C. (2002). An activation domain in the C-terminal subunit of HCF-1 is important for transactivation by VP16 and LZIP. Proc. Natl. Acad. Sci. U.S.A. 99, 13403–13408.

Luo, Y., Hitz, B. C., Gabdank, I., Hilton, J. A., Kagda, M. S., Lam, B., Myers, Z., Sud, P., Jou, J., Lin, K., et al. (2020). New developments on the Encyclopedia of DNA Elements (ENCODE) data portal. Nucleic Acids Res 48, D882–D889.

Mangone, M., Myers, M. P. and Herr, W. (2010). Role of the HCF-1 basic region in sustaining cell proliferation. PLoS ONE 5, e9020.

Mazars, R., Gonzalez-de-Peredo, A., Cayrol, C., Lavigne, A.-C., Vogel, J. L., Ortega, N., Lacroix, C., Gautier, V., Huet, G., Ray, A., et al. (2010). The THAP-zinc finger protein THAP1 associates with coactivator HCF-1 and O-GlcNAc transferase: a link between DYT6 and DYT3 dystonias. J. Biol. Chem. 285, 13364–13371.

Minocha, S. and Herr, W. (2019). Cortical and Commissural Defects Upon HCF-1 Loss in Nkx2.1-Derived Embryonic Neurons and Glia. Dev Neurobiol 79, 578–595.

Minocha, S., Sung, T.-L., Villeneuve, D., Lammers, F. and Herr, W. (2016a). Compensatory embryonic response to allele-specific inactivation of the murine X-linked gene Hcfc1. Developmental Biology 412, 1–17.

Minocha, S., Bessonnard, S., Sung, T.-L., Moret, C., Constam, D. B. and Herr, W. (2016b). Epiblast-specific loss of HCF-1 leads to failure in anterior-posterior axis specification. Dev. Biol. 418, 75–88.

Parenti, I., Rabaneda, L. G., Schoen, H. and Novarino, G. (2020). Neurodevelopmental Disorders: From Genetics to Functional Pathways. Trends in Neurosciences 43, 608–621.

Piton, A., Gauthier, J., Hamdan, F. F., Lafrenière, R. G., Yang, Y., Henrion, E., Laurent, S., Noreau, A., Thibodeau, P., Karemera, L., et al. (2011). Systematic resequencing of X-chromosome synaptic genes in autism spectrum disorder and schizophrenia. Molecular Psychiatry 16, 867–880.

Piton, A., Redin, C. and Mandel, J.-L. (2013). XLID-Causing Mutations and Associated Genes Challenged in Light of Data From Large-Scale Human Exome Sequencing. Am. J. Hum. Genet. 93, 368–383.

Quintana, A. M., Liu, F., O’Rourke, J. P. and Ness, S. A. (2011). Identification and regulation of c-Myb target genes in MCF-7 cells. BMC Cancer 11, 30.

Quintana, A. M., Geiger, E. A., Achilly, N., Rosenblatt, D. S., Maclean, K. N., Stabler, S. P., Artinger, K. B., Appel, B. and Shaikh, T. H. (2014). Hcfc1b, a zebrafish ortholog of HCFC1, regulates craniofacial development by modulating mmachc expression. Dev. Biol. 396, 94–106.

Quintana, A. M., Yu, H.-C., Brebner, A., Pupavac, M., Geiger, E. A., Watson, A., Castro, V. L., Cheung, W., Chen, S.-H., Watkins, D., et al. (2017). Mutations in THAP11 cause an inborn error of cobalamin metabolism and developmental abnormalities. Human Molecular Genetics 26, 2838–2849.

Rouillard, A. D., Gundersen, G. W., Fernandez, N. F., Wang, Z., Monteiro, C. D., McDermott, M. G. and Ma’ayan, A. (2016). The harmonizome: a collection of processed datasets gathered to serve and mine knowledge about genes and proteins. Database (Oxford) 2016, baw100.

Scalais, E., Osterheld, E., Weitzel, C., De Meirleir, L., Mataigne, F., Martens, G., Shaikh, T. H., Coughlin, C. R., Yu, H.-C., Swanson, M., et al. (2017). X-Linked Cobalamin Disorder (HCFC1) Mimicking Nonketotic Hyperglycinemia With Increased Both Cerebrospinal Fluid Glycine and Methylmalonic Acid. Pediatr. Neurol. 71, 65–69.

Uni, M. and Kurokawa, M. (2018). Role of ASXL1 mutation in impaired hematopoiesis and cellular senescence. Oncotarget 9, 36828–36829.

Ventura, A. P., Radhakrishnan, S., Green, A., Rajaram, S. K., Allen, A. N., O’Briant, K., Schummer, M., Karlan, B., Urban, N., Tewari, M., et al. (2010). Activation of the MEK – S6 pathway in high-grade ovarian cancers. Appl Immunohistochem Mol Morphol 18, 499–508.

Youn, H. S., Kim, T.-Y., Park, U.-H., Moon, S.-T., An, S.-J., Lee, Y.-K., Hwang, J.-T., Kim, E.-J. and Um, S.-J. (2017). Asxl1 deficiency in embryonic fibroblasts leads to cellular senescence via impairment of the AKT-E2F pathway and Ezh2 inactivation. Sci Rep 7, 5198.

Yu, H.-C., Sloan, J. L., Scharer, G., Brebner, A., Quintana, A. M., Achilly, N. P., Manoli, I., Coughlin, C. R., 2nd, Geiger, E. A., Schneck, U., et al. (2013). An X-Linked Cobalamin Disorder Caused by Mutations in Transcriptional Coregulator HCFC1. Am. J. Hum. Genet. 93, 506–514.

