## Supplemental Figure 1 for "Missense and nonsense mutations of the zebrafish *hcfc1a* gene result in contrasting mTor and radial glial phenotypes"

Supplementary Figure 1

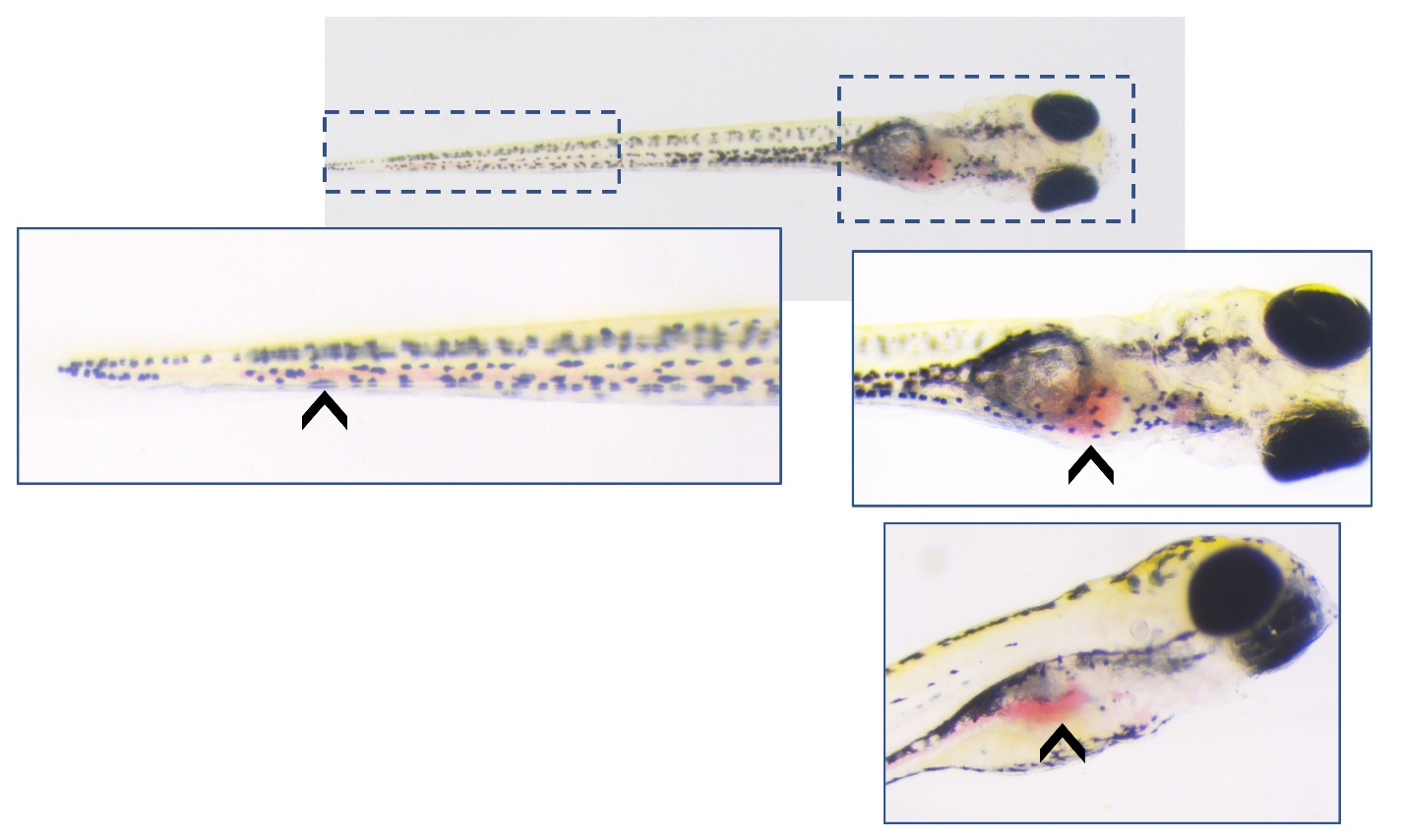

Supplementary Figure 1: Blood cell abnormalities are observed after over expression of Asxl1. Asxl1 encoding mRNA was injected at the single cell stage and embryos were grown under normal conditions until 4 days post fertilization. Red blood cell phenotypes were visualized by brightfield microscopy using a Zeiss Stereo Microscope. Arrows indicate blood pooling in specific regions of the larvae.
